# Genome wide sequencing provides evidence of adaptation to heterogeneous environments for the ancient relictual *Circaeaster agrestis* (Circaeasteraceae, Ranunculales)

**DOI:** 10.1101/2020.01.14.902643

**Authors:** Xu Zhang, Yanxia Sun, Jacob B. Landis, Jianwen Zhang, Linsen Yang, Nan Lin, Huajie Zhang, Rui Guo, Lijuan Li, Yonghong Zhang, Tao Deng, Hang Sun, Hengchang Wang

## Abstract

- Investigating the interaction between environmental heterogeneity and local adaptation is critical to understand the evolutionary history of a species, providing the premise for studying the response of organisms to rapid climate change. However, for most species how exactly the spatial heterogeneity promotes population divergence and how genomic variations contribute to adaptive evolution remain poorly understood.
- We examine the contributions of geographical and environmental variables to population divergence of the relictual, alpine herb *Circaeaster agrestis*, as well as genetic basis of local adaptation using RAD-seq and plastome data.
- We detected significant genetic structure with an extraordinary disequilibrium of genetic diversity among regions, and signals of isolation-by-distance along with isolation-by-resistance. The populations were estimated to begin diverging in the late Miocene, along with a possible ancestral distribution of the Hengduan Mountains and adjacent regions. Both environmental gradient and redundancy analyses revealed significant association between genetic variation and temperature variables. Genome-environment association analyses identified 16 putatively adaptive loci related to biotic and abiotic stress resistance.
- Our genome wide data provide new insights into the important role of environmental heterogeneity in shaping genetic structure, and access the footprints of local adaptation in an ancient relictual species, informing conservation efforts.

## Introduction

A primary goal of phylogeography is understanding how geographical and ecological factors shape the genetic structure and evolutionary history of species. Environmental factors can act in concert and vary in space, resulting in divergent natural selection leading to genetic divergence during adaptation to heterogeneous environments (Savolainen *et al.*, 2007). Such genetic divergence along environmental gradients or across varied ecological habitats, can be indicative of local adaptation, a mechanism beneficial to the long-term persistence of populations and intensifying genetic differentiation among populations (Kawecki & Ebert, 2004; Conover *et al.*, 2009; Colautti & Barrett, 2013; Savolainen *et al.*, 2013; Lowry *et al.*, 2019). Understanding the genetic basis of local adaptation will provide insights into the response of organisms to ongoing climate change, aiding in conservation efforts (Aitken *et al.*, 2008; Ahrens *et al.*, 2019).

Divergent natural selection is the driving force of genetic divergence (Darwin, 1859; Coyne & Orr, 2004), while gene flow can buffer genetic differentiation (Spieth, 1974; Goicoechea *et al.*, 2019). Evolution resulting from local adaptation depends on both the strength of local selective pressures and the homogenizing effect of connectivity among different localities. Historically, isolation by distance (IBD) (Wright 1946) have been widely employed in studies of genetic differentiation among populations. Mcrae (2006) proposed the isolation-by-resistance (IBR) model, which predicts genetic connectivity among populations in a complex landscape and provides a flexible and efficient tool to account for spatial heterogeneity, improving the understanding of environmental effect on genetic structuring. Genetic drift may reinforce the establishment and maintenance of genetic differentiation, particularly in populations with small effective size (Wright, 1946; Kimura & Crow, 1964). Quantifying the relative contributions among natural selection, genetic drift and population connectivity to genetic divergence is crucial for understanding the role of local adaptation in adaptive evolution of a species (Savolainen *et al.*, 2007; Friis *et al.*, 2018).

Accessing the molecular basis of local adaptation and identifying selective drivers is still challenging for species with limited genomic resources (Mayol *et al*., 2019). Nonetheless, with advancements in sequencing, it is now possible to study thousands of loci improving our understanding of genome-wide effects of accumulating genetic divergence and genomic properties that influence the process of local adaptation (McCormack *et al.*, 2013; Ellegren, 2014; Seehausen *et al.*, 2014; Weigel & Nordborg, 2015). In particular, highly divergent loci identified by Genome-Environment Association (GEA) analyses can be interpreted as potential targets of divergent selection associated with population-specific environmental covariables (Coop *et al.*, 2010; Rellstab *et al.*, 2015; Hoban *et al.*, 2016; Forester *et al.*, 2018). However, for most species, such as relictual alpine herbs, how exactly the spatial heterogeneity promotes population diversification and how genomic variations contribute to adaptive evolution, in particular their interaction, remains poorly understood.

In the present study, we focus on investigating the evolutionary history and local adaptation of *Circaeaster agrestis* Maxim, an annual alpine herb with a relatively narrow habitat (Fu & Bartholomew, 2001; Sun *et al.*, 2017). *Circaeaster* Maxim., with its only alliance *Kingdonia* Balf.f. & W.W. Smith, constitute the early-diverging eudicot family Circaeasteraceae (Ranunculales) (The Angiosperm Phylogeny et al., 2016). Due to special morphological characters such as open dichotomous leaf venation, this genus is of great interest to many botanists (Foster, 1971; Ren *et al.*, 2003). *Circaeaster* exhibits low morphological diversity with only one species, garnering a status of critical endangerment (Wild Plants Under State Protection in China). The distributions of *C. agrestis* is confined to the Qinghai-Tibetan Plateau (QTP) and adjacent areas which exhibit a wide range of elevations from 2,100-5,000 m. The uplift of the QTP has created extraordinarily geomorphological and climatic diversity (Mulch & Chamberlain, 2006). Due to the high mountain barrier formed by the Himalayas in the south, the central and western QTP (i.e. the Himalayas) are characterized by a cold and dry climate. In contrast, the eastern QTP (i.e. the Hengduan Mountains) and adjacent areas are associated with deep valleys and characterized mainly by a warm and wet climate (Song *et al*., 2010; Tang *et al*., 2013; Lu & Guo, 2014; Favre *et al.*, 2015). Given the high degree of geographical and ecological heterogeneity, the QTP comprises a promising model system for studying genetic signatures of local adaptation.

Mountainous regions like the QTP are often centers of endemism and species diversity hotpots (Myers et al., 2000; Noroozi *et al*., 2018) due to uplift-driven diversification (Hughes & Atchison, 2015; Chen *et al*., 2019). The uplift of the QTP produced diverse habitats facilitating rapid population divergence, but also created geographical barriers leading to isolation and fragmentation of populations (He & Jiang, 2014; Deng *et al*., 2020), the main drivers of species extinction (Hughes, 2017). Therefore, investigating genetic structure and local adaptation of species with a relatively narrow habitat will act as a proxy to comprehensively understand the role of geographical and environmental heterogeneity, while also serving as a pioneer in addressing the issues of small patchy diversity in biodiversity hotspots, aiding conservation and management efforts for threatened species.

Specifically, we generated and analyzed two types of genomic datasets: single nucleotide polymorphisms (SNPs) derived from the restriction site associated DNA sequencing (RAD-Seq) of 18 *C. agrestis* populations and 20 plastome sequences of *C. agrestis* populations with other Ranunculales representatives. We hypothesized that environmental heterogeneity may exert strong selective pressures on *C. agrestis*, driving population divergence and generating genetic variation. The aim of the present study is to illustrate the evolutionary patterns of population divergence interacting with environmental variables and access the potential genetic basis of local adaptation. Here, we raise the following questions: 1) how alpine environments affect genetic structure within species and drive population divergence, and 2) how genetic variation is associated with local adaptation? Answering these questions will help us understand species adaptation to rapidly changing climate and help to develop sound conservation programs.

## Materials and Methods

### Sampling, library preparation and sequencing

A total of 139 individuals of *C. agrestis* from 18 localities were sampled (Table S1). Sampling locations were chosen based on existing occurrence records covering the geographic and climatic distribution of the species (Fig. S1). Our field collection followed the ethics and legality of the local government. All voucher specimens were deposited in the Herbarium of Wuhan Botanical Garden (HIB) (Table S1). Total DNA was extracted from silica gel-dried leaves with a modified CTAB (Cetyl trimethylammonium bromide) method (Yang *et al.*, 2014). For RAD library construction and sequencing, genomic DNA was digested with the restriction enzyme EcoRI in a 30 ul reaction followed by ligation of P1 adapter by T4 ligase. Fragments were pooled, randomly sheared, and size-selected to 350–550 bp. A second adapter (P2) was then ligated. The ligation products were purified and PCR amplified, followed by gel purification, and size selection to 350-550 bp. Agilent 2100 Bioanaylzer and qPCR were used to qualify and quantify the library. Paired-end sequencing was performed on two lanes of Illumina HiSeq 2000 at BGI-Shenzhen (Shenzhen, Guangdong, China).

For plastome sequencing, 20 samples representing all 18 localities were chosen. For each sample, a 500-bp DNA TruSeq Illumina (Illumina Inc., San Diego, CA, USA) sequencing library was constructed using 2.5-5.0 ng sonicated DNA as input. The libraries were quantified using an Agilent 2100 Bioanalyzer (Agilent Technologies, Santa Clara, CA, USA) and real-time quantitative PCR. Libraries were multiplexed and sequenced using a 2×125 bp run on one lane of Illumina HiSeq 2000 at BGI-Shenzhen (Shenzhen, Guangdong, China).

### Processing of Illumina data

For RAD-sequencing, Illumina reads were processed into RAD-tags using the STACKS v.2.30 software pipeline (Catchen *et al.*, 2013). Samples were initially demultiplexed and filtered with PROCESS_RADTAGS. Reads with an average Phred score of at least 30, an unambiguous barcode, and restriction cut site were retained. The Perl wrapper DENOVO_MAP.PL was used for executing USTACKS, CSTACKS, and SSTACKS (Catchen *et al.*, 2013). Following the suggested protocol (Rochette & Catchen, 2017), we investigated a range of parameter values with *M* and *n* values ranging from 1 to 9 (fixing *M* = *n*) and *m* = 3. We plotted the number of polymorphic loci shared across samples (the r80 loci) and the distribution of the number of SNPs per locus.

The POPULATIONS module in the STACKS pipeline was used to produce datasets for downstream population genetic analyses. Polymorphic RAD loci that were present in all 139 individuals were retained. To validate the influence of missing data, we also employed a filtering parameter of - *r* = 0.8, in which at least 80% of individuals in a population were required to process a locus. Potential homologs were excluded by removing loci showing heterozygosity > 0.5. We further filtered our dataset with a minor allele frequency (MAF) > 0.01 and kept only biallelic SNPs. Finally, we filtered our dataset by selecting the most informative SNP based on the number of minor alleles.

For genome-skimming sequencing, raw sequence reads were filtered using Trimmomatic v.0.36 (Bolger *et al.*, 2014) by removing duplicate reads and adapter-contaminated reads. Remaining reads were directly mapped to the plastome of *C. agrestis* (NCBI accession number: KY908400; plastome size: 151,033 bp) using Geneious v9.0.2 (Kearse *et al.*, 2012). After mapping, the mean depth of coverage was 325.61x (median: 323.55x), ranging from 91.81x (WLG) to 804.08x (ZF). Initial annotations were implemented in the Plastid Genome Annotator (PGA) (Qu *et al.*, 2019), and then refined by manual correction in Geneious. The annotated plastomes were deposited in GenBank (accession numbers MT228704-MT228722).

### Population structure and genetic diversity

Population genetic structure was estimated using Bayesian clustering and principal coordinate analysis (PCoA). Bayesian clustering was performed in STRUCTURE v2.3.4 (Pritchard *et al.*, 2000) with the admixture model. As STRUCTURE analysis assumes that loci are unlinked, we thus filtered out linked SNPs using the *–write_single_snp* option in POPULATIONS (Catchen *et al.*, 2013), which exports only the first SNP per locus for analysis. To determine the optimal number of groups (*K*), we ran STRUCTURE 10 times for *K* = 1 to K = 10. Each run was performed for 200,000 Markov Chain Monte Carlo (MCMC) generations with a burn-in period of 100,000 generations. The optimal *K* was chosen using both delta-*K* method implemented in STRUCTURE HARVESTER (Earl & vonHoldt, 2012), and cross-entropy criterion implemented in *LEA* v2.8.0 (*snmf* function) (https://bioconductor.org/packages/LEA/) (Frichot. & François, 2015). The *snmf* function estimates an entropy criterion that evaluates the quality of fit of the statistical model to the data using a cross□validation technique to choose the number of ancestral populations that best explains the genotypic data (Frichot *et al*., 2014). The coefficient for cluster membership of each individual was averaged across the 10 independent runs using CLUMPP (Jakobsson & Rosenberg, 2007) and plotted using DISTRUCT (Rosenberg, 2004). We performed PCoA analyses using *dartR* v1.1.11 (*gl.pcoa* function). We tested the overall population structure using the G-statistic test (Goudet *et al.*, 1996) by *gstat.randtest* function in *adegenet* v2.1.1 (Jombart, 2008). An analysis of molecular variance (AMOVA) was performed to quantify genetic differentiation among and within populations and genetic groups using Arlequin v3.5 (Excoffier *et al.*, 2007) with significance tests of variance based on 10,000 permutations.

We estimated genetic diversity indices including nucleotide diversity (*π*), expected heterozygosity (*H*_*e*_), observed heterozygosity (*H*_*o*_) and inbreeding coefficients (*F*_IS_) using POPULATIONS. The polymorphic sites were exported in *Phylip* format and used for calculating haplotypes in DnaSP v6.12 (Rozas *et al.*, 2017). Population and group level *F*_ST_ using Weir and Cockerham’s method (Weir & Cockerham, 1984) was done using *hierfstat* (Goudet, 2005) in R v3.6.1 (R Team, 2014). The absolute differentiation (*D*_XY_) among genetic groups were measured using a Perl script provided by Ru *et al*., (2018) (Li *et al*., 2020). Finally, we assessed the contemporary effective population sizes (*N*_*e*_) for each population and group using NeEstimator v2.1 (Do *et al*., 2014). Estimates of *N*_*e*_ were calculated from the bias corrected (Waples, 2006) linkage disequilibrium method (Hill, 1981) with a minor allele frequency cutoff of 0.05 and 95% confidence intervals (CI) estimated by jackknifing.

### Population divergence

We adopted a two-step strategy for investigating the divergent time of *C. agrestis*. First, we used plastome sequences assembled from Ranunculales and newly sampled *C. agrestis* populations to estimate the diversification of *C. agrestis*. A total of 47 plastomes were included. The 79 protein coding regions (CDS) were extracted and aligned using MAFFT v7.313 (Katoh & Standley, 2013) and then concatenated using PhyloSuite v1.1.15 (Zhang *et al.*, 2019). The program BEAST2 v2.5.2 (Bouckaert *et al.*, 2014) was employed for molecular dating with concatenated matrix using a GTR+G+I substitution model selected by jModelTest v2.0.1 (Darriba *et al.*, 2012) under the Bayesian information criterion (BIC). The uncorrelated relaxed-clock mode and birth-death process were applied. The most recent common ancestors (TMRCA) of Ranunculales was constrained to a minimum age of 112 million years ago (Ma) according to the flower fossil assigned to *Teixeiraea lusitanica* von Balthazar, Pedersen & Friis (von Balthazar *et al*., 2005; Magallón *et al.*, 2015). We set a minimum of 72 Ma to constrain the diverging between Berberidaceae and Ranunculaceae (Anderson *et al*., 2005; Sun *et al*., 2018). Following the study of Bell et al., (2010), we modeled both fossil calibrations as an exponential distribution (Ho & Phillips, 2009) with a mean of 1 and an offset (hard bound constraint) that equaled the minimum age of the calibrations. The MCMC was ran for 100 million generations sampling every 1,000 generations. Tracer v1.7.1 (Rambaut *et al.*, 2018) was used to assess the effective sample size (ESS > 200) of each parameter. A maximum clade credibility tree was built by TreeAnnotator v2.5.2 (Rambaut & Drummond, 2010) using median node heights, with the initial 20% of trees discarded as burn-in.

Second, we estimated divergent time between haplotypes of *C. agrestis* using SNP sequences. All the parameters of BEAST2 were kept the same as above, with the exception of the TMRCA of *C. agrestis* constrained based on the previous results. We inferred ancestral distributions of *C. agrestis* using both statistical dispersal-extinction-cladogenesis (S-DEC) (Ree & Sanmartín, 2009) and statistical dispersal vicariance (S-DIVA) (Yu *et al.*, 2010) analyses. Both analyses were implemented in Reconstruct Ancestral State in Phylogenies (RASP) v4.0 (Yu *et al.*, 2015), using 50,000 pruned trees (only *C. agrestis* included) from posterior distribution generated by BEAST2. For both analyses, the number of maximum areas at each node was set to two, and other parameters were left as default. Five biogeographic regions were defined according to the floristic division of China proposed by Wu & Wu (1998) and the phylogeographic study of Lin *et al*., (2018): A, Qinling-Daba Mountains; B, North Hengduan Mountains; C, North QTP; D, South Hengduan Mountains; E, East Himalayan. To detect potential demographic expansions, temporal changes in the effective population size (*N*_*e*_) were inferred with Extended Bayesian skyline plots (EBSP) as implemented in BEAST2 (Heled & Drummond, 2008). We used the SNP sequences of 139 individuals and the default substitution model. The MCMC was set to 50 million generations sampling every 5,000 generations, and the first 20% was discarded as burn-in. Final graphs were produced with the custom R script provided by Heled (2010), with a burn-in cutoff of 20%.

### Effects of heterogeneous landscapes on genetic structure

To illustrate the effects of the heterogeneous landscapes on shaping genetic structure, we conducted Mantel tests for the presence of IBD and IBR. To avoid the potential correlation between geographical distance and resistance distance, a partial Mantel test was also conducted to test IBD and IBR by controlling the resistance and geographical distance. GenAlEx v6.5 (Peakall & Smouse, 2012) was employed to compute the pairwise geographic distance among 18 populations. Resistance distance was generated in CIRCUITSCAPE v4.0.5 based on circuit theory (McRae, 2006; McRae *et al*., 2008). Circuit resistance provide models of connectivity or resistance to predict patterns of dispersal among sites in a heterogeneous landscape (McRae, 2006; Dickson *et al.*, 2019). The current ecological niche model (ENM) was calculated in MAXENT (Phillips & Dudik, 2008). We used 19 bioclimatic variables available from WorldClim2 (Fick & Hijmans, 2017) at 30 arc□seconds resolution (Table S1), and actual evapotranspiration (AET; http://www.physicalgeography.net/fundamentals/8j.html). To avoid multicollinearity, we ran a Pearson correlation analysis to eliminate one of the variables in each pair with a correlation value higher than 0.9. A total 12 climatic layers was retained for analyses (Table S1).

We used environmental niches representing connectivity between populations as conductance grids to produce pairwise resistance distance, as high habitat suitability was assumed to have low resistance (Nowakowski *et al.*, 2015).Tests of the significance for the relationship between geographical/resistance distances and genetic distance among populations were implemented in *ade4* v1.7 (https://CRAN.R-project.org/package=ade4) using *mantel.rtest* with 9,999 permutations.

### Environmental variables and genetic structure

To estimate the contributions of environmental variables to genetic differentiation and to understand the turnover of allele frequencies along an environmental gradient, a gradient forest (GF) analysis was performed using *gradientForest* v0.1 (http://gradientforest.r-forge.r-project.org/). GF is a nonparametric, machine□learning regression tree approach that allows for exploration of nonlinear associations of spatial, environmental, and allelic variables (Gugger *et al.*, 2017; Bay *et al.*, 2018). The analysis partitions the allele frequency data at split values along the environmental gradients and defines the amount of variation explained as ‘split importance’ values (Jiang *et al.*, 2019). The overall predictor importance plot of GF shows the mean importance weighted by allele *R*^*2*^, and the cumulative plot for each predictor shows cumulative change along the environmental gradient. The split importance values are added cumulatively to produce a step-like curve, thus, areas with large steps in a row indicate significant influence on allelic change. The 12 climatic layers used in above analysis was employed as environmental variables in the GF analysis.

A redundancy analysis (RDA) was performed to understand the associations between genetic structure and environmental variables. We estimated the proportion of genetic variance in the populations that is explained by environmental variables using six important variables identified by GF (bio03: isothermality; AET: actual evapotranspiration; bio04: temperature seasonality; bio07: temperature annual range; bio12: annual precipitation and bio15: seasonality precipitation, see results). We constrained the dependent variables (individuals) by the explanatory variables (climate). The RDA analysis was performed using the *rda* function in *vegan* v2.5 (Oksanen *et al.*, 2018; http://CRAN.R-project.org/package=vegan). The *anova.cca* function was used to calculate overall significance and significance of each climate variable was calculated using 9,999 permutations.

### Genome-Environment Association (GEA)

To access the evidence of divergent selection acting on the genome and the genetic signatures of local adaptation, tests of associated outlier loci with environmental variation were carried out using BAYESCENV v1.1 (de Villemereuil & Gaggiotti, 2015), which extends the capabilities of the BAYESCAN algorithm by including a model that incorporates environmental data. BAYESCENV has been shown to be fairly robust to isolation by distance and a hierarchically structured scenario (de Villemereuil & Gaggiotti, 2015). For a comprehensive consideration of the environmental effect, the environmental covariable layer was calculated with principal component analysis (PCA) of the six important bioclimatic variables, and the first PC axis (PC1), which explained most of the variability (48.62%), was extracted as the input for BAYESCENV. We ran 20 pilot runs of 5,000 iterations and a burn□in of 50,000 iterations, ultimately obtaining 5,000 MCMC iterations for the analysis. Diagnostics of the log likelihoods and *F*_ST_ values for the 5,000 sampled iterations were checked using *coda* v0.19 (Plummer *et al.*, 2006) to confirm convergence and sample sizes of at least 2,500.

For functional annotation, the locus consensus sequences of all 6,120 SNPs were exported in *fasta* format. Candidate loci identified in GEA were annotated with Blast2Go v2 (Gotz *et al.*, 2008) by searching the non-redundant *Arabidopsis thaliana* protein database with records from the NCBI ESTs databases (Blastx). All the parameters were set as default with e-value < 1.0E-5.

## Results

### Sequence data processing

We produced 543 gigabyte (Gb) data containing 845,396,236 raw reads for 139 individuals of *C. agrestis* (Table S2). After filtering, the average number of used reads per sample was 4,542,944 (median: 4,883,567; Table S2). The depths of coverage for processed samples ranged from 5.77x (WLG-2) to 22.44x (HZ2-3), with a mean coverage of 11.99x (Table S2). We optimized the parameters for STACKS analysis and obtained the optimal parameters for our dataset to be *M* = *n* =3 (Fig. S2). The genome size of *C. agrestis* was estimated to be 1Gb using flow cytometry. After *de novo* assembling, we obtained 3,640,060 loci with mean genotyped sites per locus -144.49bp (a total of c.a. 0.51Gb), which comprises more than half of the genome. After strict filtering (*-r* = 1), we obtained 6,959 RAD loci containing 6,120 variant sites that were used for population genetic analyses. The mean genotyped sites per locus was 146.59bp (stderr: 0.03). After a looser filtering (*-r* = 0.8), we obtained 24,247 RAD loci containing 22,915 variant sites. The sizes of 20 sequenced plastomes from 18 populations of *C. agrestis* ranged from 150,979 bp (ZF) to 151,056 bp (SNJ) possessing the same 79 protein coding genes arranged in the same order (Table S3).

### Genetic diversity and structure

We detected an extraordinary difference of genetic diversity at population level, especially for nucleotide diversity (*π*), which ranged from 0.0008 (HZ2) to 0.1646 (DDL). The expected heterozygosity (*H*_e_) and observed heterozygosity (*H*_o_) were 0.0007 (HZ2) - 0.1555 (DDL) and 0.0014 (HZ2) - 0.2650 (DDL; Table 1), respectively. When a loose filtering strategy was employed, we obtained a similar genetic diversity pattern (Table S4). In total, 66 haplotypes were detected (Fig. 1; Table S5) and the haplotype diversity (*H*_d_) was 0.9798. Genetic differentiation between YL and HZ3 is the highest (*F*_ST_=0.9913), and that between ZG and DDL is the lowest (*F*_ST_=0.0640; Table S6; Fig. S3a). Estimates of *N*_*e*_ (Table S7) for each population ranged from 0.2 (SNJ, TSM, XLS, DDL) to 1.2 (ZF). Six sites (HZ2, HZ3, KMG, YL, ZG and GS) had negative estimates and infinite 95% CI, which may result from either a truly large *N*_*e*_ or the consequence of limited sampling (Do *et al*., 2014).

**Table 1.**
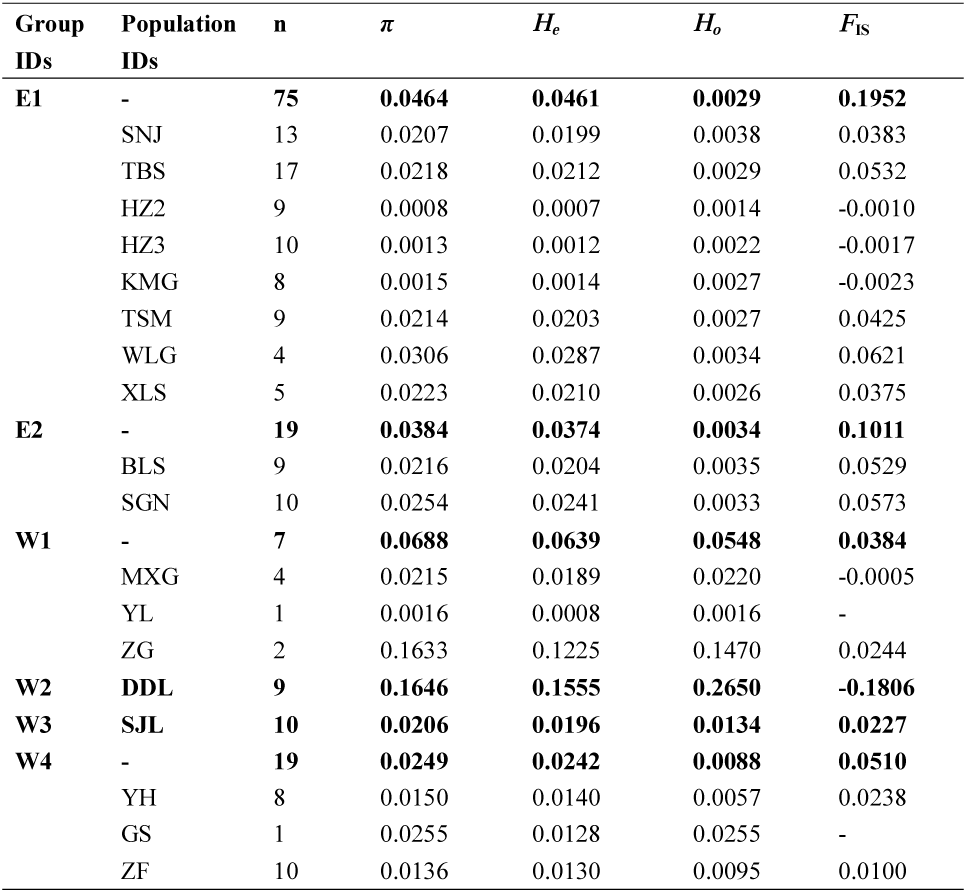
Summary of statistics calculated for the 6120 variant positions. n, number of genotype; *π*, average nucleotide diversity; *H*_*e*_, average expected heterozygosity per locus; *H*_*o*_, average observed heterozygosity per locus; the inbreeding coefficients (*F*_IS_). The statistics of six defined groups are bolded. E1, eastern group1; E2, eastern group2; W1, western group1; W2, western group2; W3, western group3; W4, western group4

**Fig. 1.**
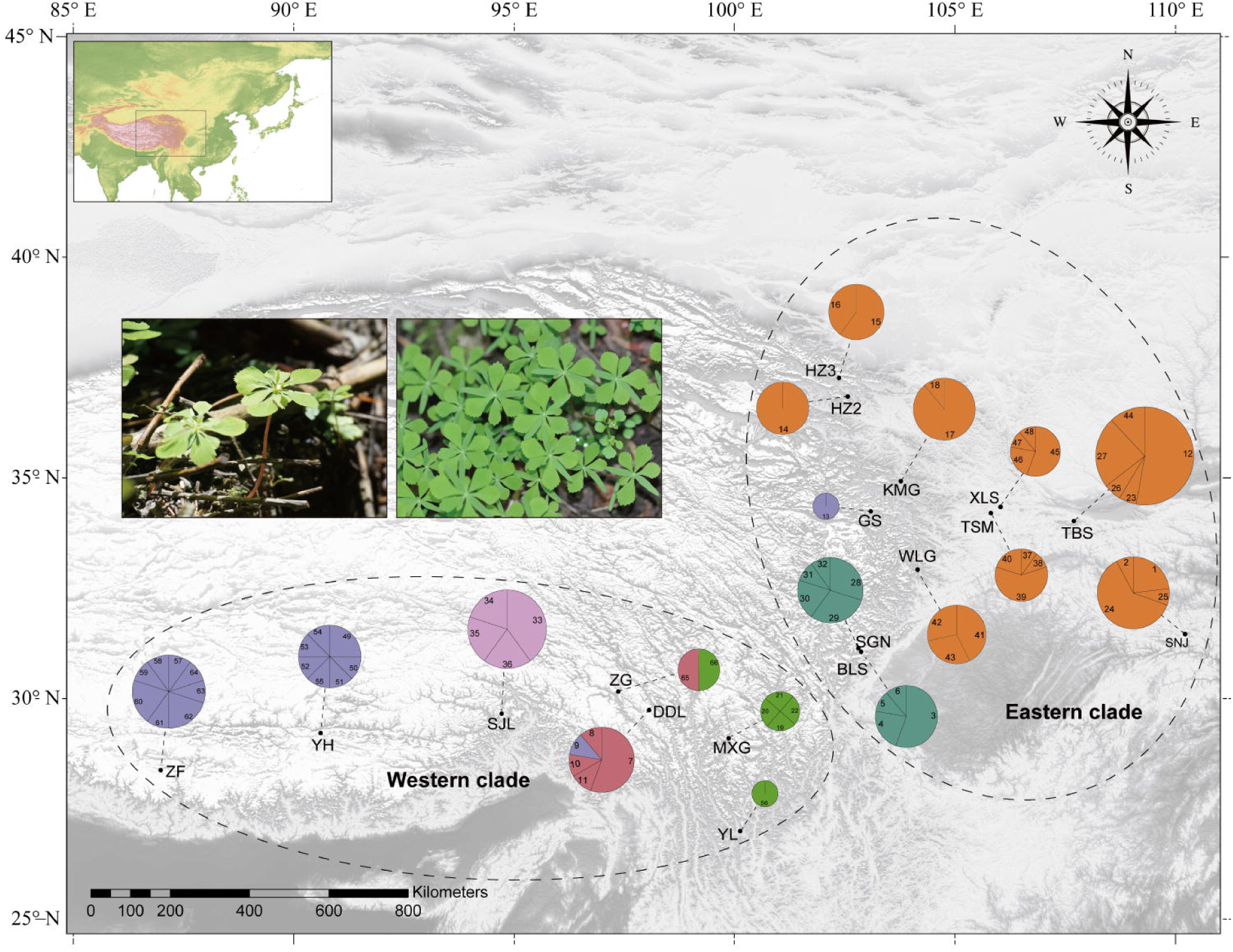
Geographical distribution of haplotypes detected in polymorphic sites of *Circaeaster agrestis* populations. Colors indicate genetic groups based on STRUCTURE and PoCA results (see also Fig. 2), with the Eastern clade being made up of the orange and dark green circles, with the Western clade consisting of the light green, light pink, purple, and blue circles. Circle sizes reflect the number of individuals sampled.

We conducted a Bayesian clustering analysis to explore the genetic structure of *C. agrestis*. The delta-*K* method identified the best-fit number (highest Δ*K* value) was two (Fig. S4a), referred to the East and West clades hereafter; the cross-entropy criterion indicated a better number of clusters was six (Fig. S4b). Considering the high *F*_ST_ among populations and the potential bias of Δ*K* method (Janes et al., 2017), we further subdivided the two clades into two and four groups (E1, E2, W1, W2, W3 and W4), respectively, which was also supported by the PoCA (Fig. 2; Fig. S5). The AMOVA revealed that 77.59% of the overall variation was distributed among six groups, with 11.27% explained by variation among populations within groups (Table S8). Within genetic groups, W2 exhibited the highest nucleotide diversity (*π*; 0.1646), while W3 had the lowest (0.0206). All genetic groups had lower observed heterozygosity than expected, with the exception of W2. In general, the East clade had lower levels of observed heterozygosity (*H*_o_, E1=0.0029, E2=0.0034, W1=0.0548, W2=0.2650, W3=0.0134, W4=0.0088) and higher levels of inbreeding coefficient (*F*_IS_, E1=0.1952, E2=0.1011, W1=0.0384, W2=-0.1806, W3=0.0227, W4=0.0510) than the West clade (Table 1). Both *F*_ST_ and *D*_XY_ showed similar patterns of divergence (Table S9 and S10; Fig. S3b), with genetic differentiation between W3 and W4 the highest (*F*_ST_ =0.8908; *D*_XY_ = 0.4307), followed by E2 and W4 (*F*_ST_=0.8451; *D*_XY_ = 0.3670), and that between E1 and E2 the lowest (*F*_ST_=0.1284; *D*_XY_ = 0.0620). Estimates of *N*_*e*_ ranged from 0.2 in W2 to 2.1 in E1 (Table S7). The G-statistic test suggested a significant population structure (*p* = 0.01**; Fig. S3c).

**Fig. 2.**
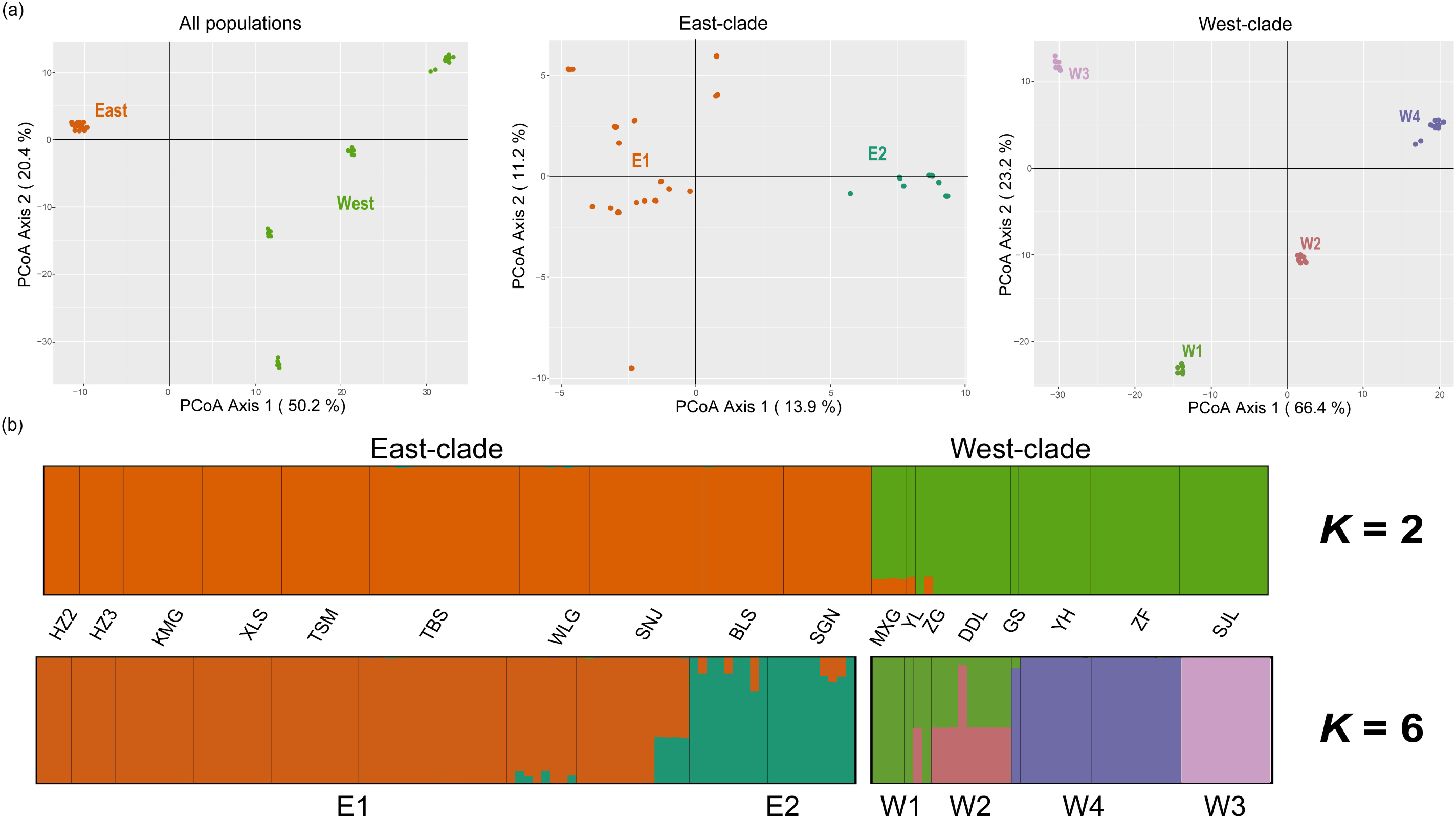
Population genetic structure in *Circaeaster agrestis* based on 6,120 SNPs from (a) PCoA for all populations, east-clade and west-clade, corresponding percentage of variation explained by each principal component is provided in **Fig. S5**; and (b) STRUCTURE with K=2 (showing the highest ΔK; Fig. S4a) and 6 (representing the divergence within east and west clades).

### Population divergence

The 47-taxa 79-CDS matrix of Ranunculales used for estimating the original divergence of *C. agrestis* was 66,117 bp in length containing 12,998 parsimony-informative sites. The divergence of *C. agrestis* from *Kingdonia* was estimated to occur in the early Eocene (stem age, ca. 52.19 Ma; 95% highest posterior density [HPD] intervals, 26.12-83.13 Ma), whereas the divergence within *C. agrestis* started in the late□Miocene (ca. 6.33 Ma; 95% HPD, 1.96-23.35 Ma) (Fig. S6). The tree topology inferred by BEAST2 using the SNP sequences identified two main clades (East and West clades) and six sub-groups (E1, E2, W1, W2, W3 and W4 groups), consistent with STRUCTURE and PoCA results. The divergence of the two main clades was estimated at appropriately 6.71 Ma (95% HPD, 5.18-7.89 Ma). The subsequent divergence between the six groups occurred mainly during the Pliocene to late Pleistocene (ca. 5.05Ma-2.90Ma) (Fig. 3a). Biogeographic analyses based on both S-DEC and S-DIVA supported Hengduan Mountains and adjacent regions as the most likely ancestral region for *C. agrestise* (Fig. 3a, 3b and Fig. S7). For the demographic inference, EBSP analysis showed no sign of population expansion or contraction in demographic history of *C. agrestis* (Fig. 3c).

**Fig. 3.**
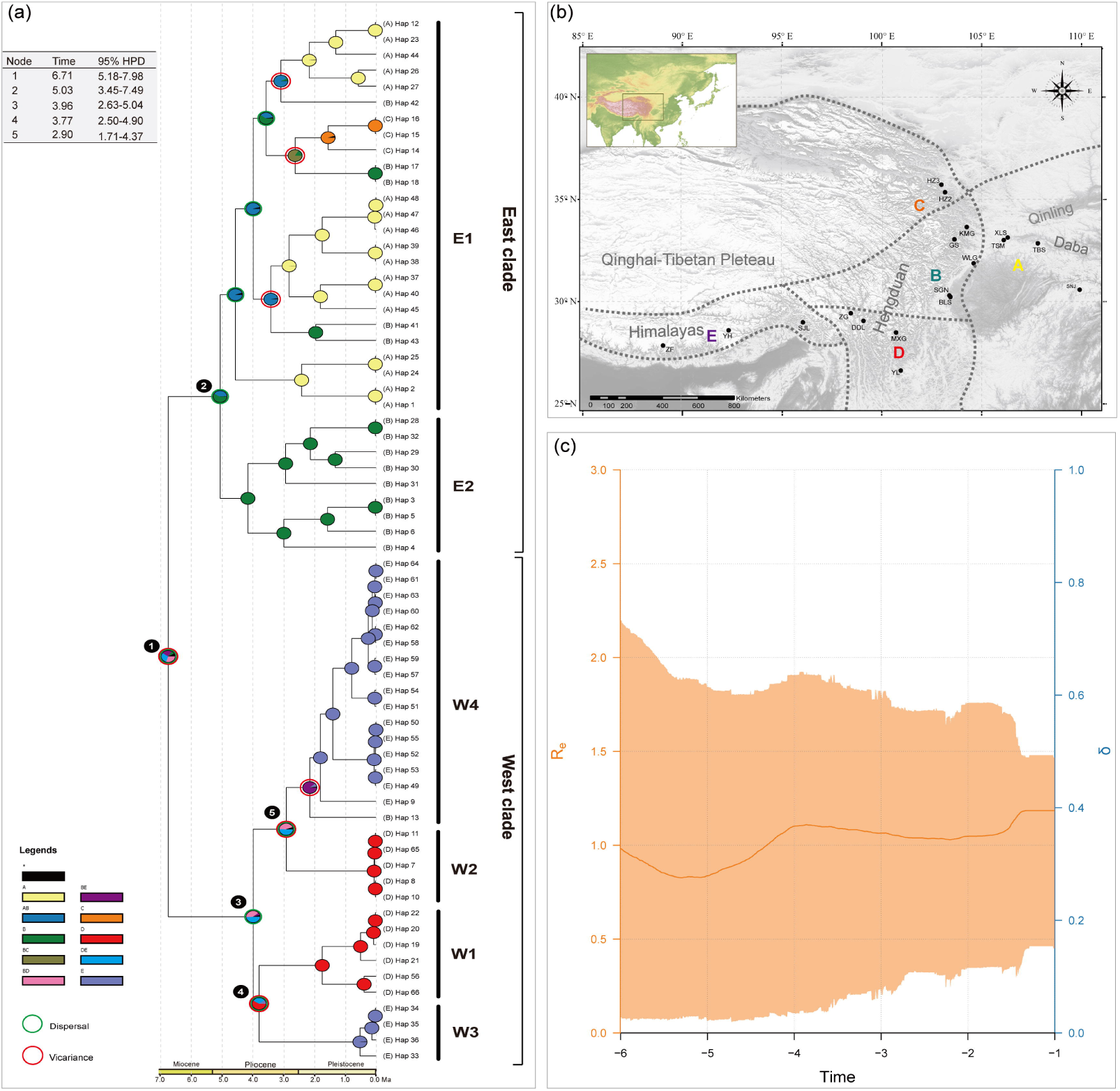
Evolutionary history of *Circaeaster agrestis*. (a) Divergent time estimation and ancestral area reconstructions of 66 haplotypes using BEAST and statistical dispersal-extinction-cladogenesis (S-DEC), respectively. Mean divergence dates and 95% HPDs for major nodes (1–5) are summarized in top left. Pie charts on each node indicate marginal probabilities for each alternative ancestral area derived from S-DEC. Results are based on a maximum area number of two. Inferred dispersal and vicariance events are indicated by green and red circle respectively. (b) Five biogeographic regions representing the current distributions of *C. agrestis*, according to the floristic division of China proposed by Wu & Wu (1998) and the study of Lin et al., (2018): A, Qinling-Daba Mountains; B, North Hengduan Mountains; C, North QTP; D, South Hengduan Mountains; E, East Himalayan regions. (c) Temporal changes in the effective population size (*N*_*e*_) changes inferred with Extended Bayesian skyline plots (EBSP) using BEAST2. Re, reproductive number, δ, death rate.

### Effects of topography and ecology

Given that the distribution range of *C. agrestis* is topographically complex, we evaluated the influence of the heterogeneous landscape in shaping genetic structure via Mantel tests. Our analyses suggested both significant patterns of IBD (*R*^2^= 0.443, *P* = 0.001**) and IBR (*R*^2^= 0.315, *P* = 0.017*) (Fig. 4a and b), which were also supported by a partial Mantel test (*R*_1_ ^2^= 0.426, *P* = 0.001**; *R*_2_ ^2^= 0.192, *P* = 0.011*). Geographical distance explained the genetic structure better than resistance distances based on current distribution of habitats. We further plotted the conductance grid derived from the IBR model to illustrate the connectivity among populations and thus predict the probabilities of successful gene flow through a complex landscape. In general, connectivity gaps among populations were observed in the western clade, reflecting increased habitat isolation in this area resulted from complex topography. In contrast, locations of eastern clade had low resistance to dispersal thus high connectivity among populations (Fig. 4c).

**Fig. 4.**
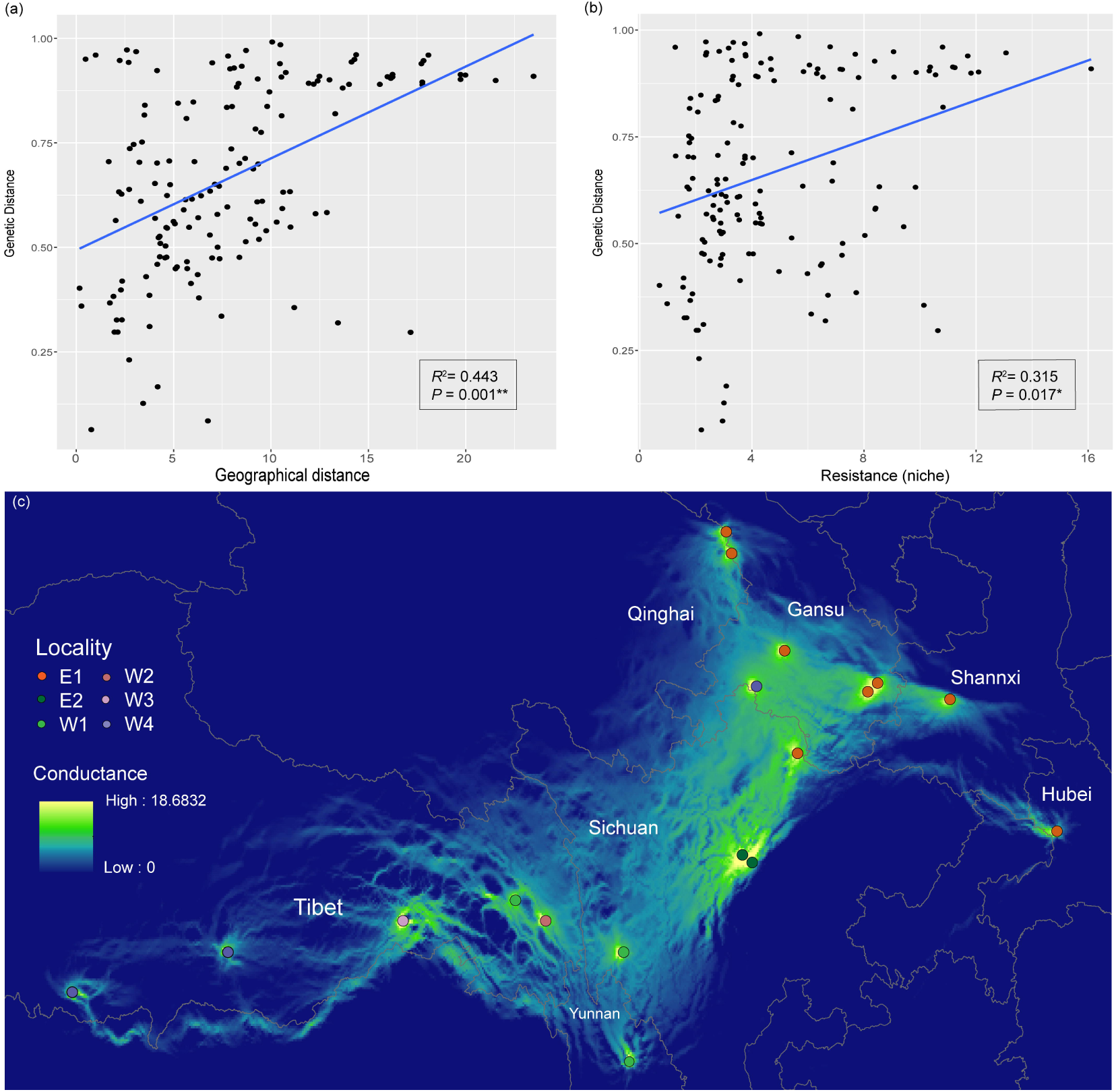
Topographic and ecological effects on genetic structure. Relationship of genetic distance and (a) geographical distance, (b) resistance distance based on climatic niche suitability, as tested by a Mantel test. (c) Conductance grid derived from the Isolation-by-Resistance (IBR) model. Grid with warm colour indicates high conductance, cold colour indicates high resistance.

### Environmental variables associated with genetic structure

Among the 12 variables used for GF analysis, isothermality (bio03) was indicated as the most important predictor. Actual evapotranspiration (AET), temperature annual range (bio07) and temperature seasonality (bio04) were also of high importance. Seasonality precipitation (bio15), and annual precipitation (bio12) showed moderate importance to allele frequencies. The other six environmental variables had lower contributions (Fig. 5a). Fig. 5b shows the cumulative change in overall allele frequencies, where changes evolved with the environmental gradient.

**Fig. 5.**
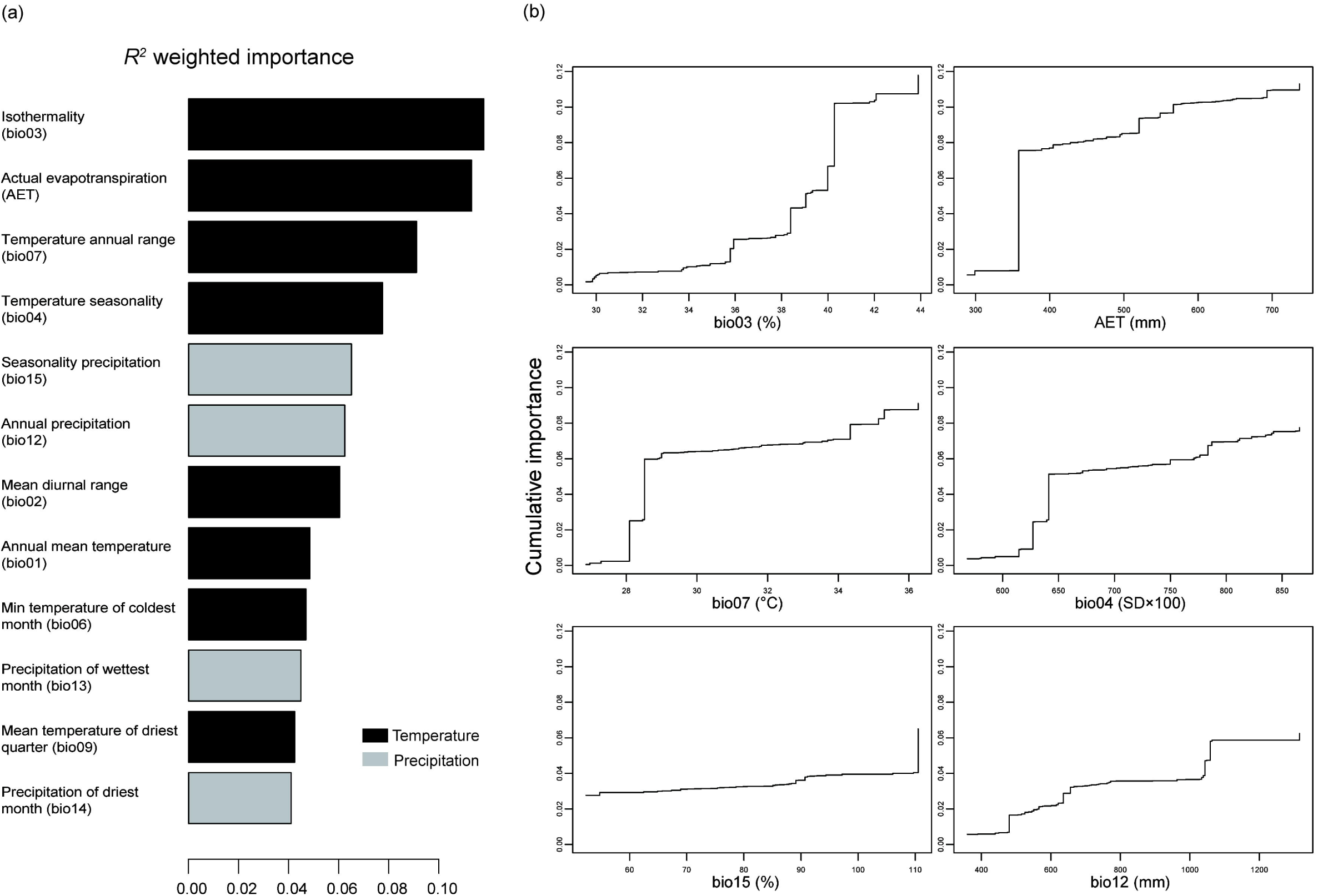
(a) R2-weighted importance of environmental variables that explain genetic gradients from gradient forest (GF) analysis. (b) Cumulative importance of allelic change along the first six environmental gradients. The units of environmental variables are provided in the parentheses. SD, standard deviation.

The RDA analysis revealed a significant amount of genetic variation among populations associated with the six important environmental variables (46.58%, *p* = 0.001). Each of the two axes explained a significant amount of variation (Axis 1: 58.55%, *P* = 0.001**; Axis 2: 27.31%, *P* = 0.001**; Fig. 6). All six environmental variables were ran separately and found to be significant. The contribution on genetic variation of each variable was generally consistent with the GF analysis. Isothermality (bio03; 18.93%, *P* = 0.001**) was the most important predictor, whereas the temperature annual range (bio07) was less important in RDA analysis (Table 2).

**Table 2.** RDA results based on six important environmental variables identified by GF analysis.

| Environmental<br>variables | Constrained<br>proportion | $F$ | $p$ |
| --- | --- | --- | --- |
| **Bio03** | 0.1893 | 31.98 | 0.001** |
| **AET** | 0.1712 | 28.31 | 0.001** |
| **Bio04** | 0.1408 | 22.45 | 0.001** |
| **Bio12** | 0.0902 | 13.59 | 0.001** |
| **Bio07** | 0.0627 | 9.17 | 0.001** |
| **Bio15** | 0.0203 | 2.84 | 0.001** |

**Fig. 6.**
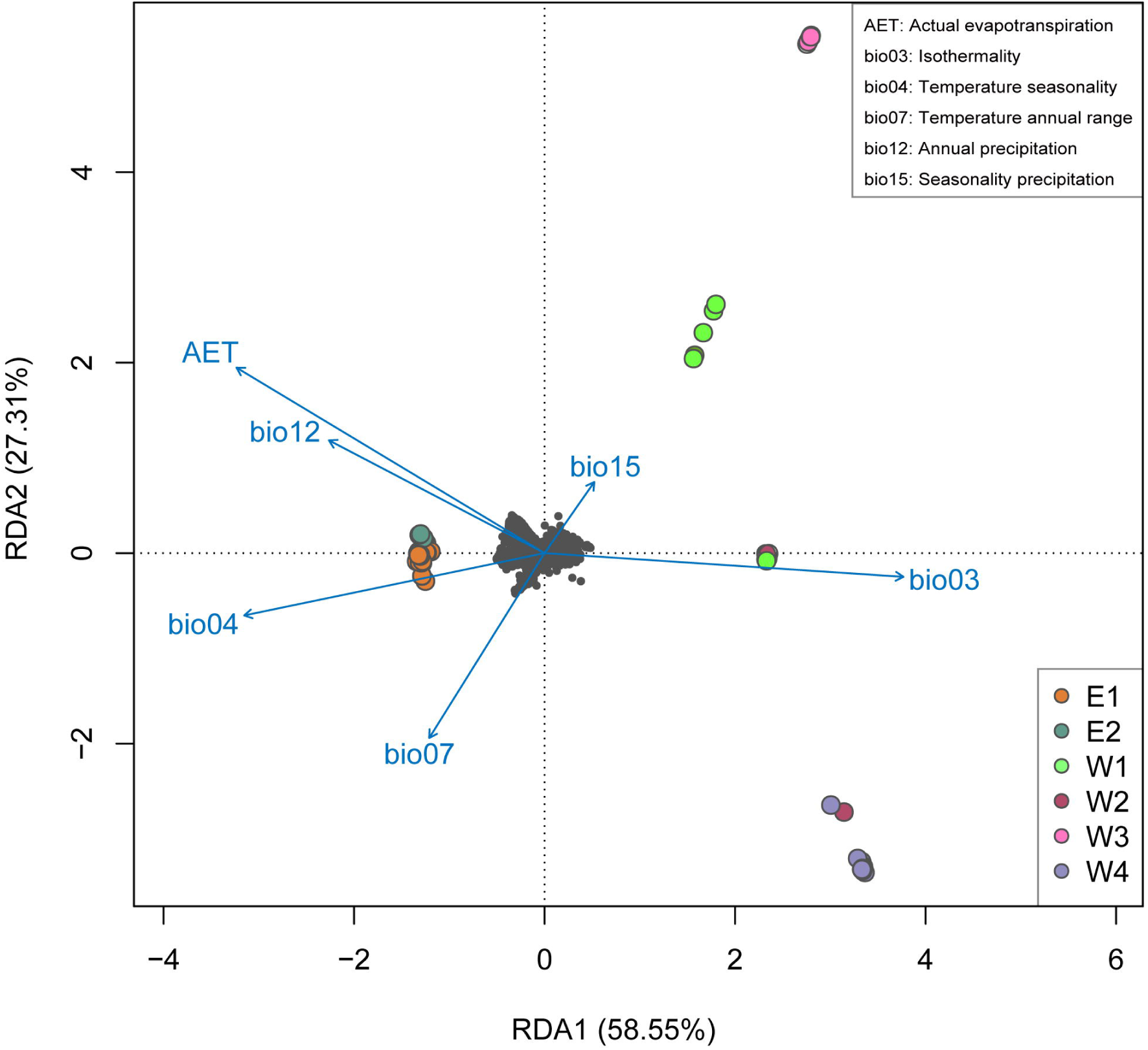
Redundancy analysis showing the relationship between the independent climate parameters and population structure. Individuals are colored points and colors represent six groups (E1, E2, W1, W2, W3, W4). Small black points are SNPs. The definition of each climate variable is provided in the top right corner.

### Genome□Environment Association (GEA)

The BAYESCENV result revealed a total of 61 loci having a significant association with environmental variables. Sixteen of the 61 loci were successfully annotated by Blast2Go with cut-off of e-value <1E-5 and a mean similarity more than 80 (Table 3; Table S11). Among these were genes related to abiotic stress response and stress-induced morphological adaptations, such as *UGT74E2* (Loci ID: 2763; Gene Ontology (GO) term: transferase activity) regulating auxin homeostasis (Tognetti *et al*., 2010), *SRK2E* (Loci ID: 8038; GO: protein kinase activity, ATP binding) involved in drought resistance (Mustilli *et al.*, 2002) and *AHK5* (Loci ID: 203056; GO: phosphorelay sensor kinase activity, phosphorelay signal transduction system) regulating stomatal state and transmitting the stress signal (Pham *et al.*, 2012), as well as genes required for the formation and development of leaves and flowers (*CYP71*, loci ID: 293398; GO: regulation of flower development, meristem structural organization leaf formation; *CSI1*, loci ID: 218068; GO: cellulose biosynthetic, pollen tube development) (Li & Luan, 2011). These loci also include genes related to oxidation-reduction process involved in cellular respiration, for example *NDUFS7* (Loci ID: 83859; GO: oxidoreductase activity, NAD binding) and *NDUFB3* (Loci ID: 219291; GO: electron transport chain).

**Table 3.**
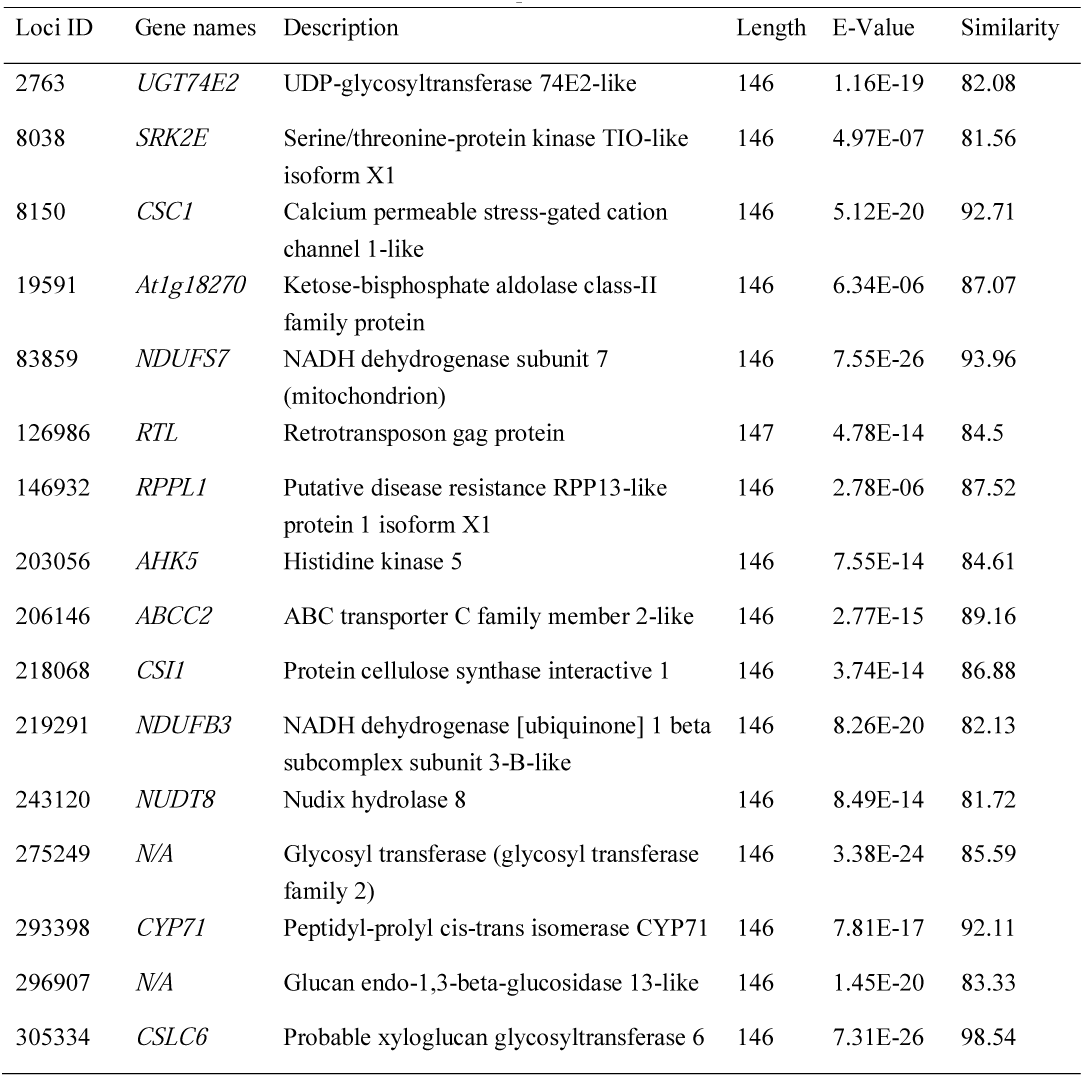
Annotation information of 16 candidate loci under selection that the E-value of blastx is less than 1E-5. Detailed functional annotation is provided in Table S11.

## Discussion

### Effects of topography on genetic structure

Our study presents the genetic structure of the palaeoendemic alpine *C. agrestis*. High haplotype diversity, uneven distribution of genetic diversity, as well as significant IBD and IBR patterns suggest restricted gene flow is likely a key factor in the high genetic differentiation among *C. agrestis* populations. The distribution area of *C. agrestis* is remarkable for its extraordinarily topographic diversity with some of the world’s most rugged mountain ranges and altitudinal gradients spanning over 5,000 m (Wen *et al.*, 2014; Favre *et al.*, 2015). As a direct consequence of the uplift of the QTP, habitat isolation induced by complex topography and frequented climatic fluctuation was invoked as an important mechanism for the high diversity of alpine plants in this area (Wen *et al.*, 2014; Sun *et al.*, 2017). Hence, dramatic terrain and climate changes probably influenced genetic connectivity and shaped current genetic structure of *C. agrestis*.

We explored the number of genetic clusters of *C. agrestis* using delta-*K*, cross-entropy criterion and PCoA analysis. Despite potential bias of delta-*K* method which exhibits a pathology toward *K*=2 (Janes et al 2017), the two distinct clades identified may reflect the early diversification of *C. agrestis*, and correspond to two unique climatic regions (Chen *et al.*, 2018). Both cross-entropy criterion and PCoA analysis indicated fine-scale structure within *C. agrestis.* For conservation purposes, fine-scale genetic structure can provide essential information for *in situ* conservation (Chung *et al.*, 2005). In particular, common garden experiments were found to be infeasible for *C. agrestis* during our field investigation, suggesting *ex situ* conservation strategies impractical.

Although previous studies suggested that drastically different levels of genetic diversity among populations may produce a bias estimate of *F*_ST_ (Charlesworth, 1998), both *F*_ST_ and *D*_XY_ showed similar pattern of genetic differentiation, revealing high isolation in the western clade. Coupled with the significant IBR pattern, habitat isolation, perhaps caused by complex topography, may be the cause for limited gene flow as well as observed high differentiation. The QTP orogeny likely triggered the vicariance among different *C. agrestis* groups, consistent with the hypothesis of diverse isolated heterogeneous habitats on sky islands (He & Jiang, 2014), which has been reported in other alpine species (e.g. *Ligularia vellerea*, Asteraceae, Yang et al., 2012; *Salix brachista*, Salicaceae, Chen *et al*., 2019; *Ficus tikoua*, Moraceae, Deng *et al*., 2020). The north-south mountain chains and valleys may provide corridors for flora exchange between the north and south but represent barriers to migration between the east and west (Chen *et al.*, 2018); likely resulting in the observed meridional differentiation of *C. agrestis* in the western clade as seen in previous studies (e.g., Gao *et al.*, 2007; Li *et al.*, 2011). Notably, the populations located in the Hengduan Mountains, exhibit high levels of genetic diversity (Table 1), which has been previously observed in plant species, such as *Parasyncalathium souliei* (Asteraceae) (Lin et al., 2018), *Taxus wallichiana* (Taxaceae) (Liu et al., 2013) and *Quercus aquifolioides* (Fagaceae) (Du et al., 2017). As the QTP blocks the cold and dry air (Liu *et al.*, 2013), the Hengduan Mountains are characterized by a much warmer and wetter climate, providing a comparatively stable environment. Although the western populations are likely to possess higher genetic diversity, they are also likely to be more vulnerable to shifting disturbance regimes and environmental changes due to lack of genetic connectivity resulted from habitats isolation. Compared to the western clade, north-south mountain barriers among different latitudes may be driving the divergence among the groups in the eastern clade. A conductance grid derived from the IBR model showed low resistance to dispersal, thus high connectivity, among populations in the eastern clade. In addition, E1 and E2 exhibit a relatively low level of genetic differentiation. This phenomenon is likely due to the similar local environments among different populations resulting from the quite moderate topographical and climatic characteristics in this region (Chen *et al.*, 2018).

The concept of effective size is key to conservation genetics, as it summarizes population status regarding inbreeding and genetic drift and provides the prospects for the sustainability of the population (Wang *et al.*, 2016). Although *N*_*e*_ is frequently less than the census population size, small values of *N*_*e*_ combined with low heterozygosity suggest high levels of inbreeding in populations of *C. agrestis*. The reduction of genetic diversity, as well as the fixation of mildly deleterious alleles that reduce reproductive fitness, influences long□term population persistence and poses a risk of extinction (Kramer & Havens, 2009; Oakley & Winn, 2012; Poudel *et al.*, 2014). In addition, episodes of low population size have a disproportionate effect on the overall value of *N*_*e*_ (Charlesworth, 2009). Thus, low *N*_*e*_ may also be attributed to the disturbance caused by human activities or possible bottleneck events in the evolutionary history of *C. agrestis.*

### Impact of environmental heterogeneity in population divergence

Our molecular dating based on plastome sequences suggested an early Eocene origin of *C. agrestis*, which is accordant with previous estimates (Ruiz-Sanchez *et al.*, 2012) and recent results from nuclear data (Sun *et al.*, 2020). The diversification of *C. agrestis* has increased since the Pliocene (Fig. 3), which is likely related to intense uplift of the Hengduan Mountains during the late Miocene to late Pliocene (Mulch & Chamberlain, 2006; Sun *et al.*, 2011; Favre *et al.*, 2015). The environmental and climatic fluctuations triggered by mountain uplifts can always lead to divergent selection and species adaptation associated with dramatic ecological niche changes (Ren *et al.*, 2017). Orogenic activities not only provide numerous new habitats favorable for diversification, but also promote population divergence due to hindered gene flow. Our biogeographic analysis revealed that the demographic evolution of *C. agrestis* has not simply been influenced by the orogeny at a particular time in history, but involves repeated episodes of dispersal and subsequent vicariance events. Coupled with the evidence of significant IBD and IBR patterns, we speculate natural selection driven by extraordinary environmental heterogeneity and geomorphological dynamics may have profound impact in the diversification of *C. agrestis*. In addition, during our field work we found the census population sizes of *C. agrestis* in the Himalaya-Hengduan Mountains to be much smaller than other sites. Previous studies have indicated a significant negative effect of small population size on adaptive evolution (Nevado *et al.*, 2019). Hence, we speculate that natural selection may play double-edged roles in the evolutionary history of any given species. On one hand, heterogeneous environments could provide more new ecological niches conducive for speciation, on the other hand, it may put the populations with weak adaptive ability to new environment in danger of extinction. Furthermore, significant IBR pattern suggest that genetic structure would be better explained by combining habitat isolation between populations than just geographical distance alone. Thus, habitat changes may have large effect on genetic structure for *C. agrestis*, indicating its sensitivity to environmental niche.

Studying local adaptation contributes to understanding the ability of populations to sustain or adapt to rapid climate change (Jia *et al.*, 2019). We associated environmental variables with genetic structure to demonstrate the impact of environmental heterogeneity on population divergence. Specifically, we found a significant association between genetic variation and temperature variables (Fig. 4; bio03, AET, bio04, bio07), suggesting temperature is an important driver of genetic variation within *C. agrestis*, as detected in previous studies of angiosperm trees (Ahrens *et al.*, 2019; Jia *et al.*, 2019; Jiang *et al.*, 2019). In fact, there is growing evidence that high-mountain environments experience more rapid changes in temperature than environments at lower elevations (Pepin *et al*., 2015; Palazzi *et al.*, 2019). Despite that, little attention has been given to understanding the mechanisms of adaptive response to climate change in high altitude areas. *C. agrestis* is habitually confined to grow under trees, shrubs or rocks, indicating sensitivity to temperature. Moreover, the vertical distribution of *C. agrestis* is extensive, ranging from 2,100 to 5,000 meters above sea level (Fu & Bartholomew, 2001), exhibiting a considerable differentiation in temperature. Therefore, it is reasonable that temperature variables play a vital role in promoting genetic divergence within *C. agrestis* and given the ongoing climate warming, temperature is likely to be a key driver for adaptation of *C. agrestis* in the future.

### Genomic signatures associated with local adaptation

BAYESCENV identified a set of genes that may be involved in a continuously adaptive process. Some genes such as pivotal kinases involved in signaling pathway regulation, may induce abiotic stress responses for defense in harsh climates in the QTP. For example, genes associated with abscisic acid (ABA) signaling pathway (*SRK2EA*) can regulate numerous ABA responses, such as stomata closure, to response to high altitudes with extreme difference in temperature (Sierla *et al.*, 2018). Considering the tiny leaves of *C. agrestis*, we speculate that stomatal regulation would be a vital process for maintaining hydration and regulating temperature. As altitude increases, hypobaric hypoxia becomes the main factor interfering with life activity. Genes related to oxidation-reduction processes involved in cellular respiration, such as *NDUFS7* and *NDUFB3*, are of importance in facilitating adaptation of *C. agrestis* to high altitude areas of hypoxia. In addition, some putatively adaptive genes (*CYP71* and *CSI1*) are associated with physiological trait regulation of vegetative and reproductive organs (Gu *et al.*, 2010). Thus, morphological adaptations may be an essential alternative for *C. agrestis* to respond to environmental stress.

A recent study charactering the genome of *Kingdonia uniflora* (sister to *C. agrestis*) calculated the nucleotide substitution rate of 1.4 × 10^−9^ per site per year (Sun *et al*., 2020), which is comparatively lower than an estimate of *Arabidopsis thaliana* (7.0 × 10^−9^) (Ossowski *et al.*, 2010) and indirect estimates based on the divergence between monocots and dicots (5.8-8.1× 10^−9^) (Wolfe *et al.*, 1987). The low nucleotide substitution rate may be evidence of low speciation rate, which likely contributes to the paucity of species in the family and the low morphological diversity of *C. agrestis* and *K. uniflora*. Low evolution rate is also indicative of low mutation rate, reducing the standing genetic variation which is the genetic source of adaptation (Lai et al., 2019). In isolated populations with high homozygosity and little standing genetic variation, only large effect alleles can escape the effects of drift becoming the targets of selection (Orr, 1998; Rausher & Delph, 2015; Sella & Barton, 2019). Additionally, many methods for detecting adaptive genetic variation often only have the power to detect loci and alleles with large phenotypic effects (Wellenreuther & Hansson, 2016; Luikart et al., 2018). Nonetheless, genetic patterns that confer adaptation to environmental heterogeneity are mostly polygenic and controlled by numerous genetic variants of small□effect (Savolainen *et al.*, 2013; Ahrens *et al.*, 2019). Therefore, the genes identified as putatively adaptive in *C. agrestis* should constitute the first step in studying genetic mechanism of local adaptation in alpine environments. Further work characterizing the whole genome is needed to obtain more precise functions of genes under selective pressure and illuminate patterns of potential polygenic adaptation (Mayol *et al*., 2019). Generally, these patterns can serve as a proxy for understanding and monitoring the adaptive process in rapidly changing environment (Ahrens *et al.*, 2019), while informing future conservation management.

## Conclusions

In this study, we demonstrate two major advances. First, we provide new insights into the genetic structure and evolutionary history of a relictual alpine herb surviving in heterogeneous environment, which can inform further conservation. Our results shed light on vital roles of mountain uplift in promoting population diversification and the dual effect of environmental heterogeneity on species evolutionary history. Second, our analyses provide a roadmap to study evolution of local adaptation of non-model species for which common garden experiments are infeasible and genomic resources are limited. By conducting multiple analyses, we associated environmental variables with genetic variation and explicitly measured the relative influence of bioclimate on population divergence. In turn, the important environmental predictors were implemented in identifying potential loci that may under divergent selection to gain in-depth knowledge of genetic basis involved in local adaptation. More importantly, our results improve our understanding of drivers and mechanisms of adaptive evolution, aiding efforts on developing sound conservation programs in facing of rapid changing climate.

## Acknowledgments

We are grateful to Jie Cai for help with sample collection in Tibet. We thank Jialiang Li and Kangshan Mao for help with the calculation of absolute differentiation (*D*_XY_). This work was supported by the Strategic Priority Research Program of Chinese Academy of Sciences (XDA20050203), the Programme Foundation for the Backbone of Scientific Research by Wuhan Botanical Garden, Chinese Academy of Sciences (Y855241G01), the Major Program of National Natural Science Foundation of China (31590823), and the National Key R and D Program of China (2017YFC0505200).

## Author contributions

HCW, HS, and YXS developed the idea and designed the experiment. XZ, JWZ, LSY, NL, HJZ, RG, LJL, YHZ and TD collected the leaf materials. XZ and YXS performed the statistical analyses; XZ, YXS, JBL, HS and HCW interpreted the results and wrote the manuscript. All authors read, edited and approved the final manuscript. XZ and YXS contributed equally to this work.

## Data accessibility

All newly sequenced plastomes were deposited in National Center for Biotechnology Information (NCBI) with accession numbers MT228704-MT228722. RAD raw read data are stored at the NCBI Sequence Read Archive in the Bioproject PRJNA616150. Genomic data sets and R scripts used in this study are available in the DRYAD archives under accession doi: 10.5061/dryad.4f4qrfj7p.

## Supporting information

**Fig. S1** The known distribution area of the Circaeaster agrestis based on all recorded sampling points available in Chinese Virtual Herbarium (CVH; http://www.cvh.ac.cn/). The green triangles indicate sampling locations in this study. The red cross indicates a distribution record in reported in 1997, but not available now based on one recent field investigation in 2018.

**Fig. S2** (a) The number of 80% polymorphic loci shared across most samples (the r80 loci) as M=n increases. (b) The distribution of the number of SNPs per locus for a range of M=n values.

**Fig. S3** (a) *F*_ST_ values within populations estimated by *hierfstat*, Group names of populations were labeled in the top of boxes. (b) *F*_ST_ values within genetic groups estimated by *hierfstat.* (c) The results of G-statistic test.

**Fig. S4** The optimal *K* value identified using (a) delta-*K* method in STRUCTURE HARVESTER, and (b) cross-entropy criterion in R package *LEA.*

**Fig. S5** Scree plot of the percentage of variation explained by each principal component (PC), corresponding to **Fig. 2a** for all populations, East-clade and West-clade, respectively.

**Fig. S6** Divergence times of Ranunculales estimated by BEAST2 with a relaxed molecular clock based on the combined protein-coding region sequences. A and B indicate fossil calibration points. C and D indicate the origination and diversification of Circaeaster agrestis, respectively. Median ages of nodes are shown with bars indicating the 95% highest posterior density intervals for each node.

**Fig. S7** Ancestral area reconstructions of 66 haplotypes using statistical dispersal vicariance (S-DIVA) analysis. Pie charts on each node indicate marginal probabilities for each alternative ancestral area derived from S-DIVA. Results are based on a maximum area number of two. Inferred dispersal and vicariance events are indicated by green and red circle respectively. Defined biogeographic regions are the same as the statistical dispersal-extinction-cladogenesis analysis (Fig. 3).

**Table S1** Summary of population information of *Circaeaster agrestis*, including the number of individuals genotyped at each localities (n), geographic information,voucher specimens information, and 20 bioclimate values (bio01-bio19 and AET) of each location. AET, the actual evapotranspiration. The units of environmental variables are provided in the parentheses. SD, standard deviation.

**Table S2** Numbers of reads of RAD-seq for each individual of *Circaeaster agrestis*.

**Table S3** Assemble and annotation information of newly sequenced plastomes of *Circaeaster agrestis* populations.

**Table S4** Summary of statistics calculated for the 22915 variant positions. n, number of genotype; π, average nucleotide diversity; He, average expected heterozygosity per locus; Ho, average observed heterozygosity per locus; the inbreeding coefficients (FIS). The statistics of six defined groups are bolded. E1, eastern group1; E2, eastern group2; W1, western group1; W2, western group2; W3, western group3; W4, western group4.

**Table S5** Haplotypes information detected in 18 populations based on 6120 SNPs.

**Table S6** Paired population-level FST estimated by *hierfstat* using Weir and Cockerham’s method.

**Table S7** Estimated Ne values using linkage disequilibrium method, implemented in NeEstimator, with a minor allele frequency cutoff of 0.05 and 95% confidence intervals (CI) estimated by jackknifing. The statistics of six defined groups are bolded. E1, eastern group1; E2, eastern group2; W1, western group1; W2, western group2; W3, western group3; W4, western group4; n, number of genotype.

**Table S8** The analysis of molecular variance (AMOVA) for SNP data among six genetic groups (E1, E2, W1, W2, W3 and W4)

**Table S9** Paired group-level FST estimated by *hierfstat* using Weir and Cockerham’s method.

**Table S10** Paired mean absolute differentiation (*D*_XY_) among genetic groups using a Perl script provided by Ru et al., (2018).

**Table S11** Detailed annotation information of sixteen genes under potential divergent selection. GO: Gene Ontology

