## Supplementary material for "Genome wide sequencing provides evidence of adaptation to heterogeneous environments for the ancient relictual *Circaeaster agrestis* (Circaeasteraceae, Ranunculales)": Figure S1-S7

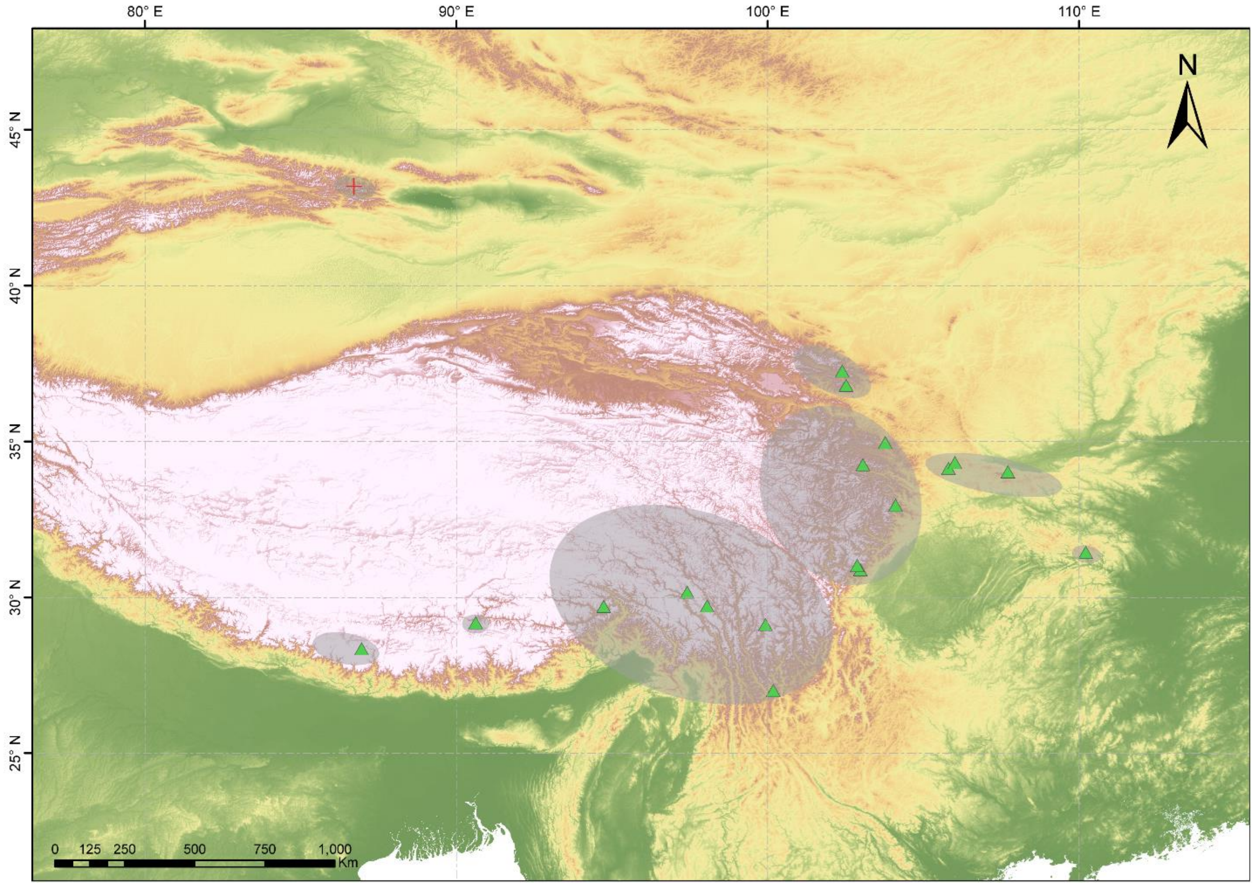


**Fig. S1** The known distribution area of the *Circaeaster agrestis* based on all recorded sampling points available in Chinese Virtual Herbarium (CVH; http://www.cvh.ac.cn/). The green triangles indicate sampling locations in this study. The red cross indicates a distribution record in reported in 1997, but not available now based on one recent field investigation in 2018.

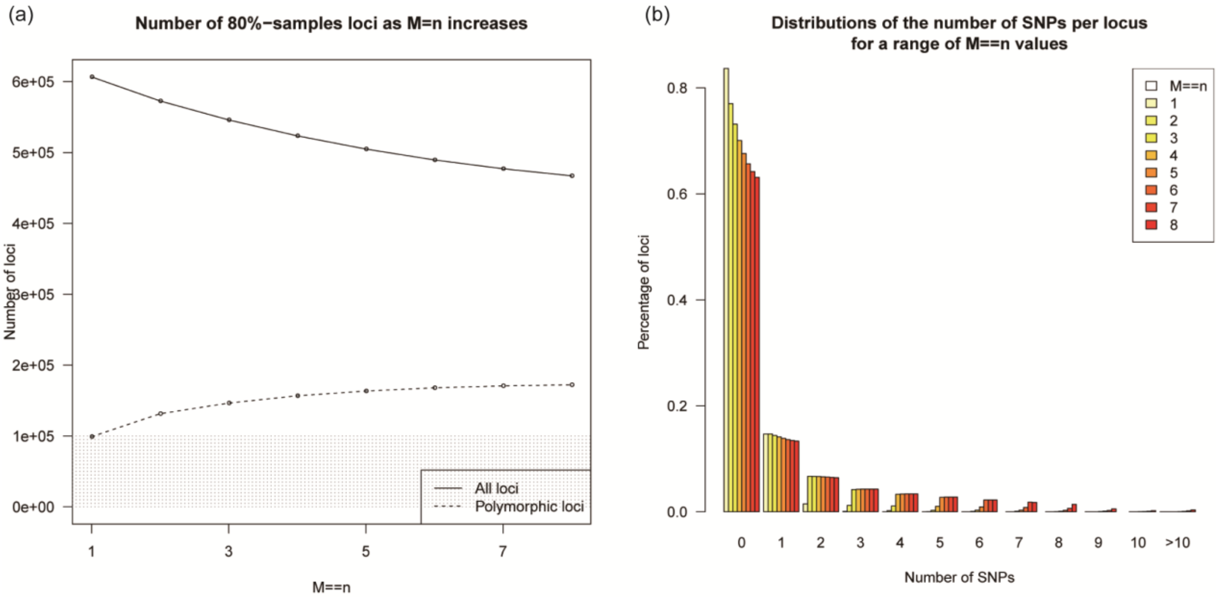


**Fig. S2** (a) The number of 80% polymorphic loci shared across most samples (the r80 loci) as M=n increases. (b) The distribution of the number of SNPs per locus for a range of M=n values.

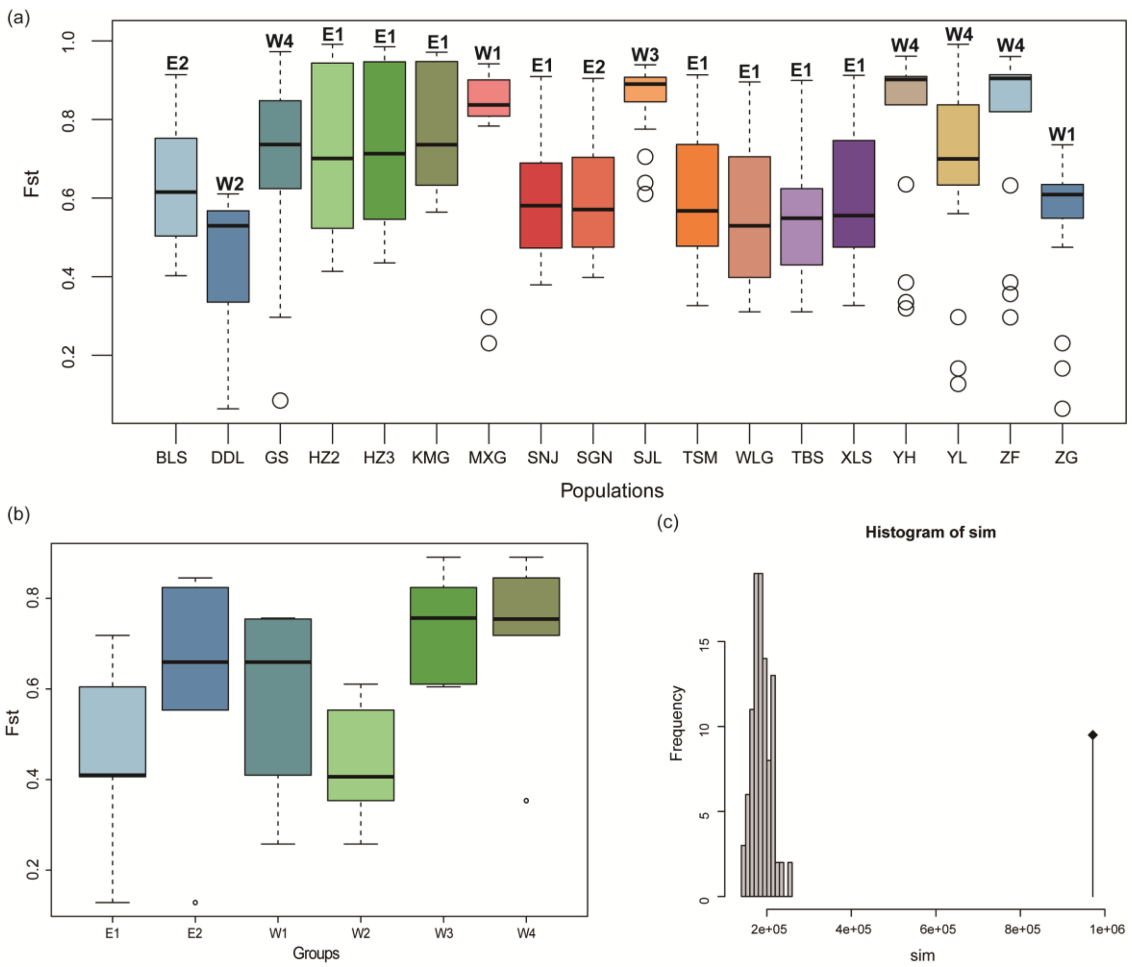


**Fig. S3** (a) $F_{ST}$ values within populations estimated by *hierfstat*, Group names of populations were labeled in the top of boxes. (b) $F_{ST}$ values within genetic groups estimated by *hierfstat*. (c) The results of G-statistic test.

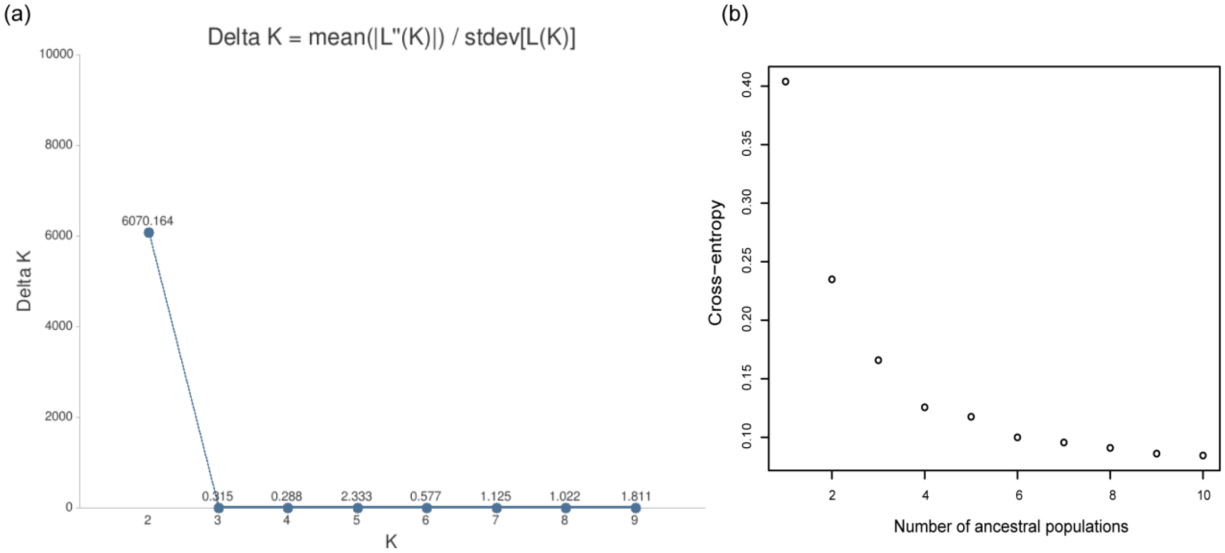


**Fig. S4** The optimal *K* value identified using (a) delta-*K* method in STRUCTURE HARVESTER, and (b) cross-entropy criterion in R package *LEA*.

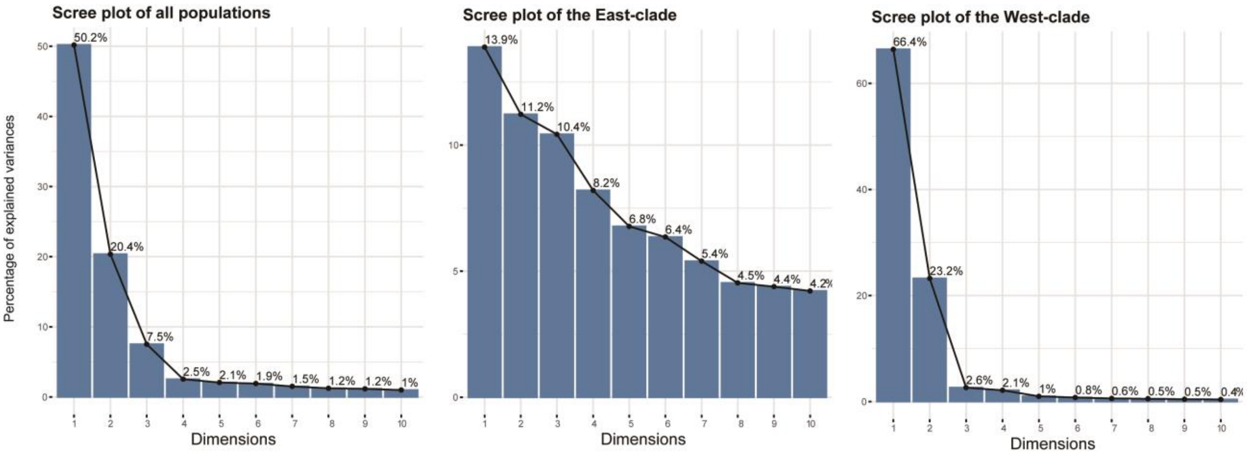


**Fig. S5** Scree plot of the percentage of variation explained by each principal component (PC), corresponding to **Fig. 2a** for all populations, East-clade and West-clade, respectively.

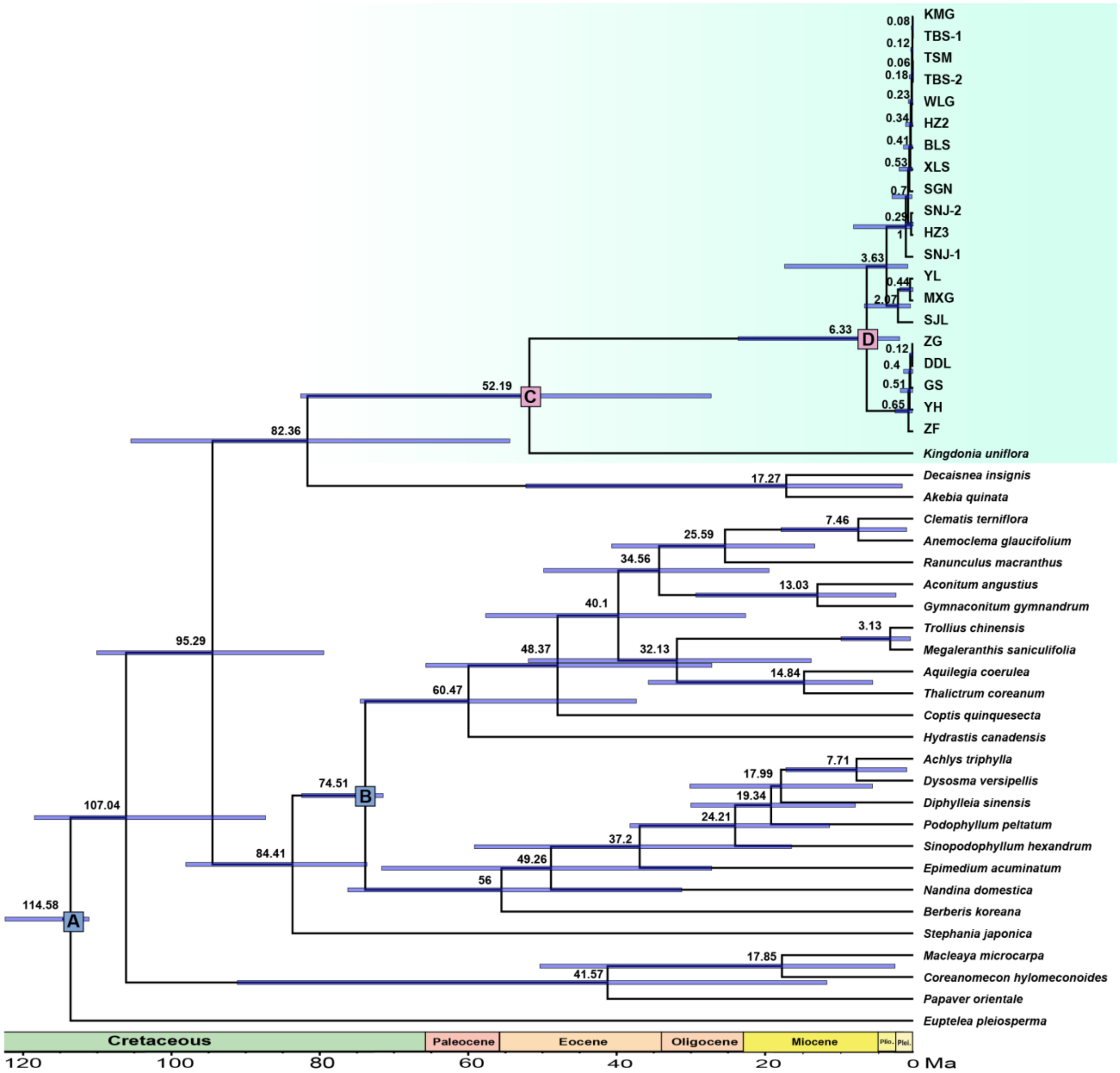


**Fig. S6** Divergence times of Ranunculales estimated by BEAST2 with a relaxed molecular clock based on the combined protein-coding region sequences. A and B indicate fossil calibration points. C and D indicate the origination and diversification of *Circaeaster agrestis*, respectively. Median ages of nodes are shown with bars indicating the 95% highest posterior density intervals for each node.

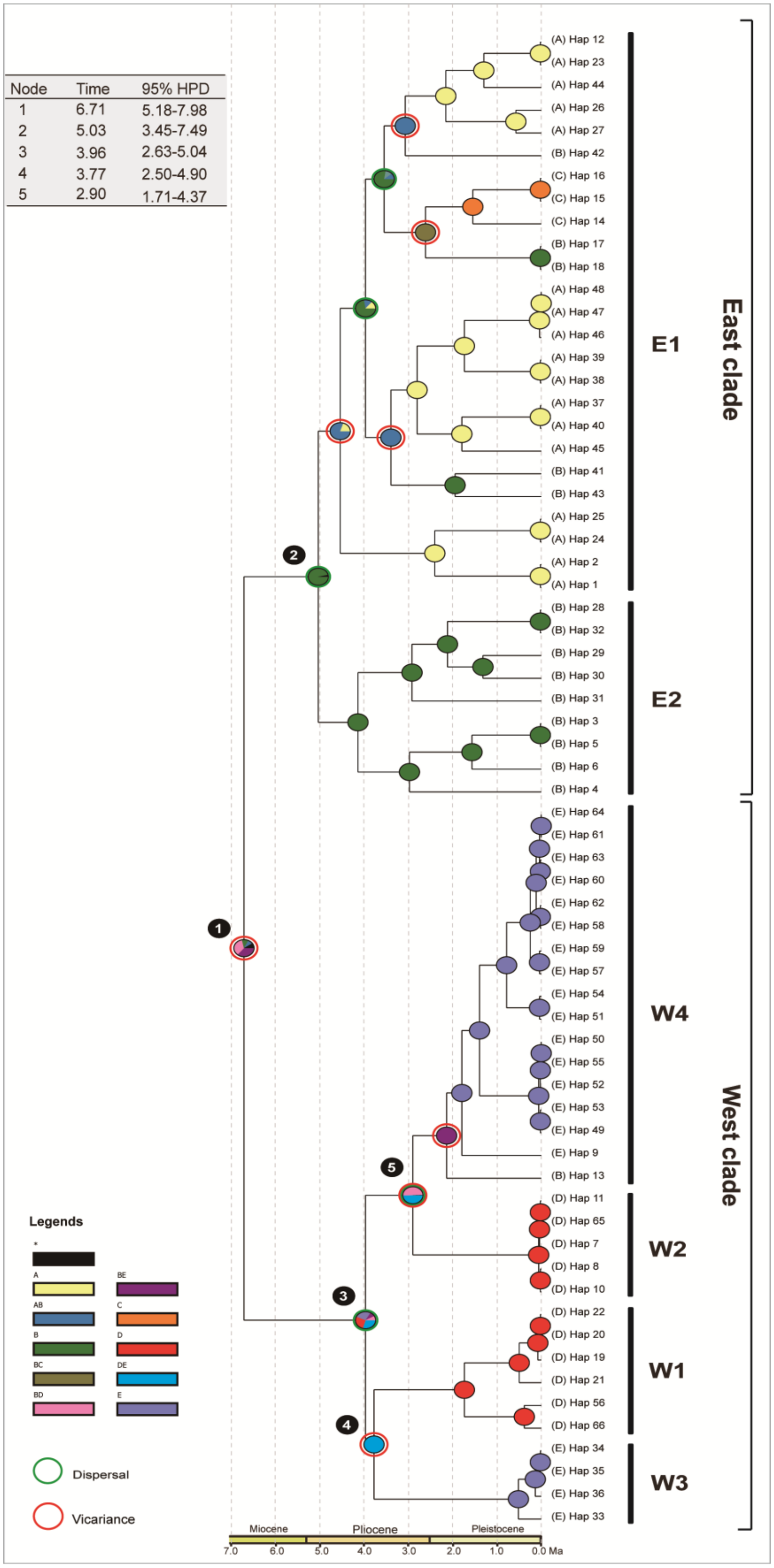


**Fig. S7** Ancestral area reconstructions of 66 haplotypes using statistical dispersal vicariance (S-DIVA) analysis. Pie charts on each node indicate marginal probabilities for each alternative ancestral area derived from S-DIVA. Results are based on a maximum area number of two. Inferred dispersal and vicariance events are indicated by green and red circle respectively. Defined biogeographic regions are the same as the statistical dispersal-extinction-cladogenesis analysis (Fig. 3).
