## Supplementary material for "Genome wide sequencing provides evidence of adaptation to heterogeneous environments for the ancient relictual *Circaeaster agrestis* (Circaeasteraceae, Ranunculales)": Table S2-S11

**Table S2** Numbers of reads of RAD-seq for each individual of *Circaeaster agrestis.*

| **Samples** |  | **Raw Reads** | **Retained Reads** | **Depths of coverage** | **BioProject ID** |
| --- | --- | --- | --- | --- | --- |
| BBY-1 |  | 6,797,509 | 5,240,364 | 14.632 | PRJNA616150 |
| BBY-3 |  | 3,705,762 | 2,818,596 | 9.269 | PRJNA616150 |
| BBY-4 |  | 5,656,881 | 4,305,940 | 12.327 | PRJNA616150 |
| BBY-5 |  | 7,906,978 | 6,317,893 | 16.884 | PRJNA616150 |
| BLS-10 |  | 3,160,616 | 2,129,059 | 7.181 | PRJNA616150 |
| BLS-2 |  | 8,670,017 | 6,456,173 | 15.611 | PRJNA616150 |
| BLS-3 |  | 7,240,775 | 5,075,737 | 11.720 | PRJNA616150 |
| BLS-4 |  | 9,654,272 | 7,699,715 | 18.737 | PRJNA616150 |
| BLS-5 |  | 6,356,124 | 4,113,099 | 9.827 | PRJNA616150 |
| BLS-6 |  | 7,669,841 | 6,079,154 | 16.263 | PRJNA616150 |
| BLS-7 |  | 6,747,987 | 5,016,195 | 12.881 | PRJNA616150 |
| BLS-8 |  | 8,184,862 | 5,821,953 | 12.463 | PRJNA616150 |
| BLS-9 |  | 6,976,031 | 5,199,821 | 13.313 | PRJNA616150 |
| DDL-1 |  | 8,668,593 | 6,915,187 | 11.925 | PRJNA616150 |
| DDL-10 |  | 5,010,292 | 3,697,062 | 7.687 | PRJNA616150 |
| DDL-2 |  | 8,620,392 | 6,548,191 | 11.165 | PRJNA616150 |
| DDL-3 |  | 4,644,115 | 3,575,296 | 10.297 | PRJNA616150 |
| DDL-4 |  | 6,494,642 | 5,064,285 | 9.044 | PRJNA616150 |
| DDL-6 |  | 6,512,476 | 4,971,664 | 9.360 | PRJNA616150 |
| DDL-7 |  | 7,840,416 | 5,958,575 | 10.351 | PRJNA616150 |
| DDL-8 |  | 5,471,296 | 4,075,921 | 8.172 | PRJNA616150 |
| DDL-9 |  | 5,409,273 | 4,087,737 | 8.316 | PRJNA616150 |
| DMG-1 |  | 8,008,479 | 6,279,341 | 16.290 | PRJNA616150 |
| DMG-2 |  | 3,478,531 | 2,425,274 | 8.173 | PRJNA616150 |
| DMG-3 |  | 5,213,529 | 4,027,197 | 11.736 | PRJNA616150 |
| DMG-4 |  | 7,951,142 | 6,160,070 | 15.860 | PRJNA616150 |
| DMG-5 |  | 5,321,548 | 3,989,369 | 11.302 | PRJNA616150 |
| GS-1 |  | 4,600,065 | 3,539,081 | 9.620 | PRJNA616150 |
| HZ2-1 |  | 6,096,807 | 4,822,775 | 13.727 | PRJNA616150 |
| HZ2-2 |  | 10,780,689 | 8,374,796 | 19.830 | PRJNA616150 |
| HZ2-3 |  | 9,494,677 | 7,945,333 | 20.437 | PRJNA616150 |
| HZ2-5 |  | 5,095,002 | 3,883,947 | 11.382 | PRJNA616150 |
| HZ3-1 |  | 7,230,520 | 5,660,905 | 14.832 | PRJNA616150 |
| HZ3-2 |  | 5,421,599 | 4,119,788 | 11.781 | PRJNA616150 |
| HZ3-3 |  | 6,193,434 | 4,951,240 | 13.317 | PRJNA616150 |
| HZ3-4 |  | 6,542,204 | 5,170,646 | 14.220 | PRJNA616150 |
| HZ3-5 |  | 6,131,511 | 4,885,777 | 13.206 | PRJNA616150 |
| KMG-11 |  | 3,759,524 | 2,778,580 | 8.784 | PRJNA616150 |
| KMG-2 |  | 8,484,240 | 6,325,503 | 15.241 | PRJNA616150 |
| KMG-3 |  | 6,240,102 | 4,627,091 | 11.922 | PRJNA616150 |
| KMG-4 |  | 5,122,138 | 3,847,691 | 10.443 | PRJNA616150 |
| KMG-5 |  | 6,845,211 | 5,334,734 | 13.904 | PRJNA616150 |
| KMG-6 |  | 7,447,075 | 5,718,570 | 14.558 | PRJNA616150 |
| KMG-7 |  | 9,107,952 | 7,257,220 | 17.737 | PRJNA616150 |
| KMG-8 |  | 7,540,056 | 5,571,231 | 13.868 | PRJNA616150 |
| KMG-9 |  | 8,095,911 | 5,846,500 | 13.078 | PRJNA616150 |
| MXG-1 |  | 3,414,560 | 2,483,675 | 8.022 | PRJNA616150 |
| MXG-2 |  | 6,432,425 | 4,453,305 | 12.167 | PRJNA616150 |
| MXG-3 |  | 8,592,316 | 6,920,778 | 18.163 | PRJNA616150 |
| MXG-4 |  | 5,821,994 | 4,453,680 | 12.168 | PRJNA616150 |
| NHG-1 |  | 6,132,438 | 4,723,232 | 13.159 | PRJNA616150 |
| NHG-2 |  | 5,738,362 | 4,552,851 | 12.894 | PRJNA616150 |
| NHG-3 |  | 7,099,202 | 5,555,108 | 15.098 | PRJNA616150 |
| NHG-4 |  | 8,614,888 | 6,669,141 | 16.766 | PRJNA616150 |
| NHG-5 |  | 7,360,568 | 5,777,548 | 15.656 | PRJNA616150 |
| NTMF-1 |  | 6,666,631 | 5,240,094 | 14.313 | PRJNA616150 |
| NTMF-3 |  | 5,780,730 | 4,504,354 | 12.630 | PRJNA616150 |
| NTMF-4 |  | 3,249,315 | 2,423,494 | 7.887 | PRJNA616150 |
| NTMF-5 |  | 5,759,922 | 4,448,420 | 12.807 | PRJNA616150 |
| NTMX-1 |  | 3,366,740 | 2,525,936 | 8.555 | PRJNA616150 |
| NTMX-2 |  | 4,652,657 | 3,643,644 | 11.015 | PRJNA616150 |
| NTMX-3 |  | 5,852,658 | 4,639,355 | 13.554 | PRJNA616150 |
| NTMX-4 |  | 5,432,390 | 4,338,567 | 12.662 | PRJNA616150 |
| NTMX-5 |  | 5,793,321 | 4,547,729 | 13.475 | PRJNA616150 |
| PAS-1 |  | 6,118,814 | 4,883,567 | 13.230 | PRJNA616150 |
| PAS-2 |  | 5,440,899 | 4,127,627 | 11.431 | PRJNA616150 |
| PAS-3 |  | 7,470,125 | 5,167,716 | 12.555 | PRJNA616150 |
| PAS-4 |  | 7,082,217 | 5,300,150 | 13.938 | PRJNA616150 |
| PAS-5 |  | 5,184,252 | 3,799,073 | 10.642 | PRJNA616150 |
| SGN-10 |  | 7,425,491 | 5,603,091 | 12.598 | PRJNA616150 |
| SGN-11 |  | 7,403,195 | 5,828,334 | 14.738 | PRJNA616150 |
| SGN-2 |  | 5,892,725 | 4,333,591 | 11.104 | PRJNA616150 |
| SGN-3 |  | 3,790,701 | 2,679,272 | 8.520 | PRJNA616150 |
| SGN-4 |  | 6,458,229 | 4,747,688 | 11.603 | PRJNA616150 |
| SGN-5 |  | 6,431,676 | 4,871,672 | 12.277 | PRJNA616150 |
| SGN-6 |  | 7,807,164 | 6,260,380 | 15.402 | PRJNA616150 |
| SGN-7 |  | 5,820,576 | 4,486,135 | 12.428 | PRJNA616150 |
| SGN-8 |  | 6,309,206 | 4,826,827 | 12.173 | PRJNA616150 |
| SGN-9 |  | 8,268,361 | 6,412,403 | 15.623 | PRJNA616150 |
| SJL-1 |  | 5,031,601 | 3,652,037 | 10.794 | PRJNA616150 |
| SJL-10 |  | 6,531,809 | 3,918,195 | 8.548 | PRJNA616150 |
| SJL-2 |  | 6,887,621 | 5,349,369 | 14.628 | PRJNA616150 |
| SJL-3 |  | 4,220,284 | 3,219,465 | 9.845 | PRJNA616150 |
| SJL-4 |  | 2,951,155 | 2,120,178 | 7.415 | PRJNA616150 |
| SJL-5 |  | 6,386,125 | 4,997,831 | 13.349 | PRJNA616150 |
| SJL-6 |  | 7,281,502 | 5,819,906 | 15.311 | PRJNA616150 |
| SJL-7 |  | 8,520,480 | 6,759,437 | 17.574 | PRJNA616150 |
| SJL-8 |  | 6,337,226 | 5,065,407 | 14.069 | PRJNA616150 |
| SJL-9 |  | 6,165,114 | 4,817,258 | 13.283 | PRJNA616150 |
| TSM-10 |  | 3,642,705 | 2,297,688 | 7.741 | PRJNA616150 |
| TSM-11 |  | 6,723,540 | 4,838,362 | 11.862 | PRJNA616150 |
| TSM-2 |  | 6,953,722 | 4,943,207 | 12.141 | PRJNA616150 |
| TSM-3 |  | 6,051,996 | 3,828,076 | 10.253 | PRJNA616150 |
| TSM-4 |  | 5,408,178 | 3,509,111 | 9.539 | PRJNA616150 |
| TSM-5 |  | 4,764,368 | 3,391,893 | 9.791 | PRJNA616150 |
| TSM-6 |  | 5,963,104 | 4,018,852 | 11.085 | PRJNA616150 |
| TSM-7 |  | 7,155,513 | 5,142,912 | 12.698 | PRJNA616150 |
| TSM-8 |  | 3,982,360 | 2,521,413 | 8.165 | PRJNA616150 |
| TSM-9 |  | 7,098,461 | 4,734,396 | 12.114 | PRJNA616150 |
| WLG-2 |  | 4,903,683 | 1,597,184 | 5.770 | PRJNA616150 |
| WLG-3 |  | 6,582,766 | 4,393,611 | 10.800 | PRJNA616150 |
| WLG-4 |  | 7,557,951 | 5,535,134 | 12.734 | PRJNA616150 |
| WLG-5 |  | 7,262,544 | 5,541,930 | 13.158 | PRJNA616150 |
| WLG-6 |  | 5,054,478 | 3,459,178 | 9.199 | PRJNA616150 |
| WLG-7 |  | 5,648,962 | 3,854,478 | 9.856 | PRJNA616150 |
| WLG-8 |  | 8,088,211 | 5,937,660 | 13.934 | PRJNA616150 |
| WLG-9 |  | 4,564,786 | 3,157,024 | 8.548 | PRJNA616150 |
| WTW-1 |  | 5,665,055 | 4,518,625 | 12.764 | PRJNA616150 |
| WTW-4 |  | 6,054,096 | 4,889,632 | 14.006 | PRJNA616150 |
| XLS-10 |  | 5,603,338 | 3,517,234 | 9.644 | PRJNA616150 |
| XLS-2 |  | 7,552,600 | 5,227,981 | 12.694 | PRJNA616150 |
| XLS-3 |  | 4,607,575 | 2,490,634 | 7.888 | PRJNA616150 |
| XLS-4 |  | 6,362,744 | 4,080,629 | 10.901 | PRJNA616150 |
| XLS-5 |  | 5,838,080 | 3,915,498 | 10.314 | PRJNA616150 |
| XLS-6 |  | 6,786,921 | 4,714,362 | 12.366 | PRJNA616150 |
| XLS-7 |  | 6,800,838 | 4,479,718 | 11.830 | PRJNA616150 |
| XLS-8 |  | 7,188,556 | 4,966,270 | 13.171 | PRJNA616150 |
| XLS-9 |  | 4,520,680 | 2,878,150 | 9.252 | PRJNA616150 |
| YH-10 |  | 4,614,240 | 3,698,897 | 11.097 | PRJNA616150 |
| YH-2 |  | 5,854,924 | 4,823,199 | 13.534 | PRJNA616150 |
| YH-3 |  | 4,299,807 | 3,573,413 | 11.227 | PRJNA616150 |
| YH-5 |  | 6,532,700 | 5,276,075 | 14.668 | PRJNA616150 |
| YH-6 |  | 3,766,329 | 2,825,203 | 8.831 | PRJNA616150 |
| YH-7 |  | 3,060,238 | 2,366,300 | 7.927 | PRJNA616150 |
| YH-8 |  | 3,778,028 | 3,050,145 | 9.571 | PRJNA616150 |
| YH-9 |  | 3,950,944 | 3,039,724 | 9.298 | PRJNA616150 |
| YL-1 |  | 5,739,304 | 4,472,093 | 12.297 | PRJNA616150 |
| ZF-1 |  | 2,105,068 | 1,325,590 | 5.798 | PRJNA616150 |
| ZF-10 |  | 4,005,061 | 2,993,168 | 9.263 | PRJNA616150 |
| ZF-2 |  | 6,054,738 | 4,786,914 | 13.425 | PRJNA616150 |
| ZF-3 |  | 4,865,204 | 3,666,230 | 10.726 | PRJNA616150 |
| ZF-4 |  | 6,640,295 | 5,146,130 | 14.108 | PRJNA616150 |
| ZF-5 |  | 5,219,256 | 3,939,509 | 11.150 | PRJNA616150 |
| ZF-6 |  | 6,085,703 | 4,730,425 | 12.924 | PRJNA616150 |
| ZF-7 |  | 3,017,083 | 2,121,199 | 7.459 | PRJNA616150 |
| ZF-8 |  | 5,120,347 | 3,806,124 | 10.969 | PRJNA616150 |
| ZF-9 |  | 4,473,441 | 3,395,806 | 9.918 | PRJNA616150 |
| ZG-1 |  | 4,633,849 | 3,169,436 | 6.747 | PRJNA616150 |
| ZG-2 |  | 6,613,105 | 5,225,024 | 14.010 | PRJNA616150 |
| Sum. |  | 845,396,236 | 631,469,242 | 1665.907 | - |
| Average |  | 6,081,987 | 4,542,944 | 11.985 | - |

**Table S3** Assemble and annotation information of newly sequenced plastomes of *Circaeaster agrestis* populations.

| **Population ID** | **Individual ID** | **Mapped reads** | **Length** | **%GC** | **Protein coding genes** | **NCBI accession number** |
| --- | --- | --- | --- | --- | --- | --- |
| BLS | BLS-3 | 176,015 | 151,042 | 38.1 | 79 | MT228704 |
| DDL | DDL-6 | 490,732 | 150,983 | 38.2 | 79 | MT228705 |
| GS | GS-1 | 297,414 | 150,995 | 38.2 | 79 | MT228707 |
| HZ2 | HZ2-1 | 237,867 | 151,053 | 38.2 | 79 | MT228708 |
| HZ3 | HZ3-4 | 145,876 | 151,073 | 38.2 | 79 | MT228709 |
| KMG | KMG-6 | 190,731 | 151,052 | 38.2 | 79 | MT228710 |
| MXG | MXG-2 | 382,353 | 151,028 | 38.1 | 79 | MT228711 |
| SGN | SGN-7 | 213,797 | 151,048 | 38.2 | 79 | MT228714 |
| SJL | SJL-1 | 419,889 | 151,041 | 38.2 | 79 | MT228715 |
| SNJ | BBY-4 | 422,698 | 151,033 | 38.2 | 79 | KY908400 |
|  | NTMF-4 | 383,020 | 151,046 | 38.2 | 79 | MT228713 |
| TBS | DMG-5 | 293,780 | 151,054 | 38.2 | 79 | MT228706 |
|  | WTW-1 | 291,881 | 151,054 | 38.1 | 79 | MT228712 |
| TSM | TSM-7 | 268,972 | 151,056 | 38.2 | 79 | MT228716 |
| WLG | WLG-2 | 92,456 | 151,054 | 38.2 | 79 | MT228717 |
| XLS | XLS-6 | 192,105 | 151,050 | 38.2 | 79 | MT228718 |
| YH | YH-9 | 578,015 | 150,998 | 38.2 | 79 | MT228719 |
| YL | YL-1 | 333,572 | 151,038 | 38.1 | 79 | MT228720 |
| ZF | ZF-1 | 809,329 | 150,979 | 38.2 | 79 | MT228721 |
| ZG | ZG-1 | 336,191 | 151,002 | 38.2 | 79 | MT228722 |

**Table S4** Summary of statistics calculated for the 22915 variant positions. n, number of genotype; π, average nucleotide diversity; He, average expected heterozygosity per locus; Ho, average observed heterozygosity per locus; the inbreeding coefficients (FIS). The statistics of six defined groups are bolded. E1, eastern group1; E2, eastern group2; W1, western group1; W2, western group2; W3, western group3; W4, western group4.

| **Group IDs** | **Population IDs** | **n** | ***π*** | ***H_e_*** | ***H_o_*** | ***F*_IS_** |
| --- | --- | --- | --- | --- | --- | --- |
| E1 | SNJ | 13 | 0.0208 | 0.0199 | 0.0043 | 0.0371 |
|  | TBS | 17 | 0.0229 | 0.0222 | 0.0046 | 0.0501 |
|  | HZ2 | 9 | 0.0014 | 0.0012 | 0.0023 | -0.0015 |
|  | HZ3 | 10 | 0.0013 | 0.0012 | 0.0022 | -0.0015 |
|  | KMG | 8 | 0.0015 | 0.0014 | 0.0025 | -0.0020 |
|  | TSM | 9 | 0.0188 | 0.0178 | 0.0028 | 0.0372 |
|  | WLG | 4 | 0.0289 | 0.0270 | 0.0032 | 0.0574 |
|  | XLS | 5 | 0.0211 | 0.0199 | 0.0025 | 0.0353 |
| E2 | BLS | 9 | 0.0215 | 0.0203 | 0.0038 | 0.0508 |
|  | SGN | 10 | 0.0231 | 0.0220 | 0.0028 | 0.0529 |
| W1 | MXG | 4 | 0.0201 | 0.0176 | 0.0201 | 0.0009 |
|  | YL | 1 | 0.0026 | 0.0013 | 0.0026 | - |
|  | ZG | 2 | 0.1946 | 0.1460 | 0.1805 | 0.0212 |
| W2 | DDL | 9 | 0.2048 | 0.1931 | 0.3258 | -0.2209 |
| W3 | SJL | 10 | 0.0170 | 0.0161 | 0.0113 | 0.0181 |
| W4 | GS | 8 | 0.0246 | 0.0123 | 0.0246 | - |
|  | YH | 1 | 0.0163 | 0.0152 | 0.0060 | 0.0262 |
|  | ZF | 10 | 0.0150 | 0.0142 | 0.0088 | 0.0155 |

**Table S5** Haplotypes information detected in 18 populations based on 6120 SNPs.

| Population | Hap. | Num. | Hap. | Num. | Hap. | Num. | Hap. | Num. | Hap. | Num. | Hap. | Num. | Hap. | Num. | Hap. | Num. | Total Hap. | Total ind. |
| --- | --- | --- | --- | --- | --- | --- | --- | --- | --- | --- | --- | --- | --- | --- | --- | --- | --- | --- |
| SNJ | H1 | 3 | H2 | 1 | H24 | 8 | H25 | 1 |  |  |  |  |  |  |  |  | 4 | 13 |
| BLS | H3 | 5 | H4 | 2 | H5 | 1 | H6 | 1 |  |  |  |  |  |  |  |  | 4 | 9 |
| DDL | H7 | 5 | H8 | 1 | H9 | 1 | H10 | 1 | H11 | 1 |  |  |  |  |  |  | 5 | 9 |
| TBS | H12 | 9 | H26 | 1 | H27 | 4 | H44 | 2 | H23 | 1 |  |  |  |  |  |  | 5 | 17 |
| GS | H13 | 1 |  |  |  |  |  |  |  |  |  |  |  |  |  |  | 1 | 1 |
| HZ2 | H14 | 4 |  |  |  |  |  |  |  |  |  |  |  |  |  |  | 1 | 4 |
| HZ3 | H15 | 3 | H16 | 2 |  |  |  |  |  |  |  |  |  |  |  |  | 2 | 5 |
| KMG | H17 | 8 | H18 | 1 |  |  |  |  |  |  |  |  |  |  |  |  | 2 | 9 |
| MXG | H19 | 1 | H20 | 1 | H21 | 1 | H22 | 1 |  |  |  |  |  |  |  |  | 4 | 4 |
| SGN | H28 | 3 | H29 | 3 | H30 | 2 | H31 | 1 | H32 | 1 |  |  |  |  |  |  | 5 | 10 |
| SJL | H33 | 2 | H34 | 1 | H35 | 6 | H36 | 1 |  |  |  |  |  |  |  |  | 4 | 10 |
| TSM | H37 | 1 | H38 | 1 | H39 | 5 | H40 | 2 |  |  |  |  |  |  |  |  | 4 | 10 |
| WLG | H41 | 3 | H42 | 2 | H43 | 2 |  |  |  |  |  |  |  |  |  |  | 3 | 8 |
| XLS | H45 | 5 | H46 | 2 | H47 | 1 | H48 | 1 |  |  |  |  |  |  |  |  | 4 | 9 |
| YH | H49 | 2 | H50 | 1 | H51 | 1 | H52 | 1 | H53 | 1 | H54 | 1 | H55 | 1 |  |  | 7 | 8 |
| YL | H56 | 1 |  |  |  |  |  |  |  |  |  |  |  |  |  |  | 1 | 1 |
| ZF | H57 | 1 | H58 | 1 | H59 | 1 | H60 | 2 | H61 | 1 | H62 | 2 | H63 | 1 | H64 | 1 | 8 | 10 |
| ZG | H65 | 1 | H66 | 1 |  |  |  |  |  |  |  |  |  |  |  |  | 2 | 2 |
| Total |  |  |  |  |  |  |  |  |  |  |  |  |  |  |  |  | 66 | 139 |

Hap., haplotypes, Num., number, ind., individual.

**Table S6** Paired population‐level *F*_ST_ estimated by *hierfstat* using Weir and Cockerham's method.

| Populations | BLS | DDL | GS | HZ2 | HZ3 | KMG | MXG | SNJ | SGN | SJL | TSM | WLG | TBS | XLS | YH | YL | ZF | ZG |
| --- | --- | --- | --- | --- | --- | --- | --- | --- | --- | --- | --- | --- | --- | --- | --- | --- | --- | --- |
| BLS | - | 0.5565 | 0.7520 | 0.6153 | 0.6234 | 0.7017 | 0.8401 | 0.5004 | 0.4023 | 0.8919 | 0.5096 | 0.4194 | 0.4656 | 0.5034 | 0.9093 | 0.7067 | 0.9136 | 0.6140 |
| DDL | 0.5565 | - | 0.0849 | 0.4762 | 0.5133 | 0.5966 | 0.2972 | 0.5806 | 0.5620 | 0.6105 | 0.5676 | 0.5294 | 0.5930 | 0.5555 | 0.3352 | 0.1270 | 0.3558 | 0.0640 |
| GS | 0.7520 | 0.0849 | - | 0.9723 | 0.9684 | 0.9599 | 0.8476 | 0.6893 | 0.7036 | 0.7751 | 0.7363 | 0.7051 | 0.6239 | 0.7464 | 0.3192 | 0.9576 | 0.2965 | 0.4747 |
| HZ2 | 0.6153 | 0.4762 | 0.9723 | - | 0.9503 | 0.9473 | 0.9290 | 0.5192 | 0.5480 | 0.9074 | 0.5232 | 0.4596 | 0.4136 | 0.5265 | 0.9434 | 0.9913 | 0.9394 | 0.7009 |
| HZ3 | 0.6234 | 0.5133 | 0.9684 | 0.9503 | - | 0.9425 | 0.9332 | 0.5397 | 0.5709 | 0.9182 | 0.5475 | 0.4764 | 0.4349 | 0.5458 | 0.9494 | 0.9847 | 0.9466 | 0.7128 |
| KMG | 0.7017 | 0.5966 | 0.9599 | 0.9473 | 0.9425 | - | 0.9415 | 0.6464 | 0.6527 | 0.9392 | 0.6327 | 0.5643 | 0.5694 | 0.6274 | 0.9607 | 0.9711 | 0.9601 | 0.7356 |
| MXG | 0.8401 | 0.2972 | 0.8476 | 0.9290 | 0.9332 | 0.9415 | - | 0.8149 | 0.8165 | 0.8448 | 0.8347 | 0.8083 | 0.7831 | 0.8369 | 0.9028 | 0.2972 | 0.9007 | 0.2307 |
| SNJ | 0.5004 | 0.5806 | 0.6893 | 0.5192 | 0.5397 | 0.6464 | 0.8149 | - | 0.4728 | 0.8900 | 0.4532 | 0.3791 | 0.4298 | 0.4489 | 0.9018 | 0.6333 | 0.9092 | 0.5833 |
| SGN | 0.4023 | 0.5620 | 0.7036 | 0.5480 | 0.5709 | 0.6527 | 0.8165 | 0.4728 | - | 0.8834 | 0.4775 | 0.3981 | 0.4493 | 0.4751 | 0.8983 | 0.6501 | 0.9044 | 0.5896 |
| SJL | 0.8919 | 0.6105 | 0.7751 | 0.9074 | 0.9182 | 0.9392 | 0.8448 | 0.8900 | 0.8834 | - | 0.8925 | 0.8720 | 0.8813 | 0.8904 | 0.9230 | 0.7050 | 0.9272 | 0.6387 |
| TSM | 0.5096 | 0.5676 | 0.7363 | 0.5232 | 0.5475 | 0.6327 | 0.8347 | 0.4532 | 0.4775 | 0.8925 | - | 0.3263 | 0.3827 | 0.3596 | 0.9082 | 0.6877 | 0.9134 | 0.6088 |
| WLG | 0.4194 | 0.5294 | 0.7051 | 0.4596 | 0.4764 | 0.5643 | 0.8083 | 0.3791 | 0.3981 | 0.8720 | 0.3263 | - | 0.3108 | 0.3265 | 0.8900 | 0.6511 | 0.8953 | 0.5787 |
| TBS | 0.4656 | 0.5930 | 0.6239 | 0.4136 | 0.4349 | 0.5694 | 0.7831 | 0.4298 | 0.4493 | 0.8813 | 0.3827 | 0.3108 | - | 0.3671 | 0.8888 | 0.5603 | 0.8992 | 0.5486 |
| XLS | 0.5034 | 0.5555 | 0.7464 | 0.5265 | 0.5458 | 0.6274 | 0.8369 | 0.4489 | 0.4751 | 0.8904 | 0.3596 | 0.3265 | 0.3671 | - | 0.9077 | 0.6996 | 0.9122 | 0.6105 |
| YH | 0.9093 | 0.3352 | 0.3192 | 0.9434 | 0.9494 | 0.9607 | 0.9028 | 0.9018 | 0.8983 | 0.9230 | 0.9082 | 0.8900 | 0.8888 | 0.9077 | - | 0.8372 | 0.3853 | 0.6347 |
| YL | 0.7067 | 0.1270 | 0.9576 | 0.9913 | 0.9847 | 0.9711 | 0.2972 | 0.6333 | 0.6501 | 0.7050 | 0.6877 | 0.6511 | 0.5603 | 0.6996 | 0.8372 | - | 0.8196 | 0.1665 |
| ZF | 0.9136 | 0.3558 | 0.2965 | 0.9394 | 0.9466 | 0.9601 | 0.9007 | 0.9092 | 0.9044 | 0.9272 | 0.9134 | 0.8953 | 0.8992 | 0.9122 | 0.3853 | 0.8196 | - | 0.6320 |
| ZG | 0.6140 | 0.0640 | 0.4747 | 0.7009 | 0.7128 | 0.7356 | 0.2307 | 0.5833 | 0.5896 | 0.6387 | 0.6088 | 0.5787 | 0.5486 | 0.6105 | 0.6347 | 0.1665 | 0.6320 | - |

**Table S7** Estimated *N_e_* values using linkage disequilibrium method, implemented in NeEstimator, with a minor allele frequency cutoff of 0.05 and 95% confidence intervals (CI) estimated by jackknifing. The statistics of six defined groups are bolded. E1, eastern group1; E2, eastern group2; W1, western group1; W2, western group2; W3, western group3; W4, western group4; n, number of genotype.

| **Group IDs** | **Population IDs** | **n** | ***N_e_*** | **Jackknife CI** | |
| --- | --- | --- | --- | --- | --- |
|  |  |  |  | **Low** | **High** |
| **E1** | **-** | **75** | **2.1** | **1.7** | **2.7** |
|  | SNJ | 13 | 0.2 | 0.2 | 0.2 |
|  | TBS | 17 | 0.4 | 0.3 | 0.6 |
|  | HZ2 | 9 | -1.4 | Infinite | Infinite |
|  | HZ3 | 10 | -1.7 | Infinite | Infinite |
|  | KMG | 8 | -3.4 | Infinite | Infinite |
|  | TSM | 9 | 0.2 | 0.2 | 0.2 |
|  | WLG | 4 | 0.5 | 0.4 | 0.5 |
|  | XLS | 5 | 0.2 | 0.2 | 0.2 |
| **E2** | **-** | **19** | **0.9** | **0.6** | **1.4** |
|  | BLS | 9 | 0.5 | 0.2 | 1.4 |
|  | SGN | 10 | 0.7 | 0.3 | 1.6 |
| **W1** | **-** | **7** | **0.3** | **0.1** | **0.9** |
|  | MXG | 4 | 0.8 | 0.2 | Infinite |
|  | YL | 1 | -0.3 | Infinite | Infinite |
|  | ZG | 2 | -1.2 | Infinite | Infinite |
| **W2** | **DDL** | **9** | **0.2** | **0** | **1.4** |
| **W3** | **SJL** | **10** | **0.8** | **0.6** | **1** |
| **W4** | **-** | **19** | **0.9** | **0.5** | **1.5** |
|  | YH | 8 | 0.6 | 0.4 | 0.9 |
|  | GS | 1 | -0.3 | Infinite | Infinite |
|  | ZF | 10 | 1.2 | 0.7 | 1.9 |

**Table S8** The analysis of molecular variance (AMOVA) for SNP data among six genetic groups (E1, E2, W1, W2, W3 and W4).

| **Source of variation** | **d.f.** | **Variance component** | **Percentage of variation (%)** | **Fixation index** |
| --- | --- | --- | --- | --- |
| Among groups | 5 | 632.68 | 77.59 | *F*_CT_ = 0.78*  *F*_SC_ = 0.50*  *F*_ST_ = 0.89* |
| Within groups | 12 | 91.92 | 11.27 |  |
| Within populations | 260 | 90.80 | 11.14 |  |

d.f., degree of freedom. Significance: *, *P* < 0.001.

**Table S9** Paired group‐level *F*_ST_ estimated by *hierfstat* using Weir and Cockerham's method.

| Groups | E1 | E2 | W1 | W2 | W3 | W4 |
| --- | --- | --- | --- | --- | --- | --- |
| E1 | - | 0.1284 | 0.4097 | 0.4061 | 0.6044 | 0.7181 |
| E2 | 0.1284 | - | 0.6592 | 0.5533 | 0.8239 | 0.8451 |
| W1 | 0.4097 | 0.6592 | - | 0.2574 | 0.7564 | 0.7543 |
| W2 | 0.4061 | 0.5533 | 0.2574 | - | 0.6105 | 0.3534 |
| W3 | 0.6044 | 0.8239 | 0.7564 | 0.6105 | - | 0.8908 |
| W4 | 0.7181 | 0.8451 | 0.7543 | 0.3534 | 0.8908 | - |

**Table S10** Paired mean absolute differentiation (*D*_XY_) among genetic groups using a Perl script provided by Ru *et al*., (2018).

| Groups | E1 | E2 | W1 | W2 | W3 | W4 |
| --- | --- | --- | --- | --- | --- | --- |
| E1 | - | 0.0620 | 0. 2668 | 0. 3076 | 0. 3494 | 0. 3646 |
| E2 | 0.0620 | - | 0. 2694 | 0. 3103 | 0. 3519 | 0. 3670 |
| W1 | 0.2668 | 0.2694 | - | 0. 1909 | 0. 2842 | 0. 3165 |
| W2 | 0.3076 | 0.3103 | 0.1909 | - | 0. 3514 | 0. 1731 |
| W3 | 0.3494 | 0.3519 | 0.2842 | 0.3514 | - | 0. 4307 |
| W4 | 0.3646 | 0.3670 | 0.3165 | 0.1731 | 0.4307 | - |

**Table S11** Detailed annotation information of sixteen genes under potential divergent selection. GO: Gene Ontology.

| Loci ID | Gene names | Description | GO IDs | GO Names |
| --- | --- | --- | --- | --- |
| 2763 | *UGT74E2* | UDP-glycosyltransferase 74E2-like | F: GO:0016758 | F: transferase activity, transferring hexose groups |
| 8038 | *SRK2E* | serine/threonine-protein kinase TIO-like isoform X1 | F: GO:0004672;  F: GO:0005524;  P: GO:0006468 | F: protein kinase activity; F: ATP binding; P: protein phosphorylation |
| 8150 | *CSC1* | calcium permeable stress-gated cation channel 1-like | C: GO:0016021 | C: integral component of membrane |
| 19591 | *At1g18270* | ketose-bisphosphate aldolase class-II family protein | F: GO:0003824;  C: GO:0005634;  P: GO:0005975;  F: GO:0008270;  F: GO:0016491;  F: GO:0016832;  F: GO:0050661;  F: GO:0051287;  P: GO:0055114 | F: catalytic activity; C: nucleus; P: carbohydrate metabolic process; F: zinc ion binding; F: oxidoreductase activity; F: aldehyde-lyase activity; F: NADP binding; F: NAD binding; P: oxidation-reduction process |
| 83859 | *NDUFS7* | NADH dehydrogenase subunit 7 (mitochondrion) | C: GO:0005739;  F: GO:0016651;  F: GO:0048038;  F: GO:0051287;  P: GO:0055114 | C: mitochondrion; F: oxidoreductase activity, acting on NAD(P)H; F: quinone binding; F: NAD binding; P: oxidation-reduction process |
| 126986 | *RTL* | Retrotransposon gag protein | F: GO:0003676;  P: GO:0015074 | F: nucleic acid binding; P: DNA integration |
| 146932 | *RPPL1* | putative disease resistance RPP13-like protein 1 isoform X1 | F: GO:0043531 | F:ADP binding |
| 203056 | *AHK5* | histidine kinase 5 | F: GO:0000155;  P: GO:0000160;  P: GO:0023014 | F: phosphorelay sensor kinase activity; P: phosphorelay signal transduction system; P: signal transduction by protein phosphorylation |
| 206146 | *ABCC2* | ABC transporter C family member 2-like | C: GO:0000325;  F: GO:0005524;  C: GO:0005774;  C: GO:0016021;  F: GO:0016887;  F: GO:0042626;  P: GO:0055085 | C: plant-type vacuole; F: ATP binding; C: vacuolar membrane; C: integral component of membrane; F: ATPase activity; F: ATPase-coupled transmembrane transporter activity; P: transmembrane transport |
| 218068 | *CSI1* | protein CELLULOSE SYNTHASE INTERACTIVE 1 | C:GO:0005794; F:GO:0008017; P:GO:0009414; C:GO:0009506; P:GO:0009833; C:GO:0009898; P:GO:0009901; P:GO:0010208; P:GO:0010215; C:GO:0010330; C:GO:0016021; F:GO:0016874; P:GO:0030244; C:GO:0036449; P:GO:0043622; P:GO:0048467; P:GO:0048868; P:GO:0051211; P:GO:0051592; C:GO:0055028; P:GO:0070507; P:GO:0072699; P:GO:2000067 | C:Golgi apparatus; F:microtubule binding; P:response to water deprivation; C:plasmodesma; P:plant-type primary cell wall biogenesis; C:cytoplasmic side of plasma membrane; P:anther dehiscence; P:pollen wall assembly; P:cellulose microfibril organization; C:cellulose synthase complex; C:integral component of membrane; F:ligase activity; P:cellulose biosynthetic process; C:microtubule minus-end; P:cortical microtubule organization; P:gynoecium development; P:pollen tube development; P:anisotropic cell growth; P:response to calcium ion; C:cortical microtubule; P:regulation of microtubule cytoskeleton organization; P:protein localization to cortical microtubule cytoskeleton; P:regulation of root morphogenesis |
| 219291 | *NDUFB3* | NADH dehydrogenase [ubiquinone] 1 beta subcomplex subunit 3-B-like | C: GO:0005747;  C: GO:0016021;  P: GO:0022900 | C: mitochondrial respiratory chain complex I; C: integral component of membrane; P: electron transport chain |
| 243120 | *NUDT8* | Nudix hydrolase 8 | F: GO:0035529;  F: GO:0047631;  F: GO:0051287 | F: NADH pyrophosphatase activity; F:ADP-ribose diphosphatase activity; F: NAD binding |
| 275249 | *N/A* | glycosyl transferase (glycosyl transferase family 2) | C: GO:0005739;  C: GO:0016021;  F: GO:0016740 | C: mitochondrion; C: integral component of membrane; F: transferase activity |
| 293398 | *CYP71* | peptidyl-prolyl cis-trans isomerase CYP71 | P:GO:0000413; F:GO:0003682; F:GO:0003755; C:GO:0005634; P:GO:0006457; P:GO:0009909; P:GO:0009933; P:GO:0010082; P:GO:0010305; P:GO:0010338; P:GO:0010358; P:GO:0031060; F:GO:0042393; P:GO:0048440; P:GO:0048443; P:GO:0048453 | P:protein peptidyl-prolyl isomerization; F:chromatin binding; F:peptidyl-prolyl cis-trans isomerase activity; C:nucleus; P:protein folding; P:regulation of flower development; P:meristem structural organization; P:regulation of root meristem growth; P:leaf vascular tissue pattern formation; P:leaf formation; P:leaf shaping; P:regulation of histone methylation; F:histone binding; P:carpel development; P:stamen development; P:sepal formation |
| 296907 | *N/A* | glucan endo-1,3-beta-glucosidase 13-like | F: GO:0042973;  P: GO:0005975;  P: GO:0071555 | F: glucan endo-1,3-beta-D-glucosidase activity; P: carbohydrate metabolic process; P: cell wall organization |
| 305334 | *CSLC6* | probable xyloglucan glycosyltransferase 6 | C: GO:0005768;  C: GO:0005802;  C: GO:0016021;  F: GO:0016740;  P: GO:0048868 | C: endosome; C: trans-Golgi network; C: integral component of membrane; F: transferase activity; P: pollen tube development |
